# Non-invasive measurement of neurotransmitter-specific glucose metabolism in the human brain using proton-observed proton-edited ^13^C-MRS (POPE^13^C-MRS)

**DOI:** 10.64898/2026.03.13.711600

**Authors:** Antoine Cherix, Oliver Haermson, Mohamed Tachrount, Jon Campbell, William T Clarke, Jason P Lerch, Charlotte Stagg

## Abstract

Non-invasive measurement of neurotransmitter-specific glucose metabolism in the human brain remains a major challenge, limiting mechanistic insight into excitatory–inhibitory imbalance across neurological and psychiatric disorders. Existing approaches either lack neurotransmitter specificity or require specialised hardware that constrains clinical applicability. Here, we introduce proton-observed proton-edited ^13^C magnetic resonance spectroscopy (POPE^13^C-MRS), an approach that enables non-invasive detection of glutamate, GABA, and lactate metabolism using standard proton MRI hardware. The method combines administration of ^13^C-labelled glucose with targeted proton editing to detect ^1^H-^13^C satellite resonances. Using a cross-species proof-of-concept approach, we demonstrate consistent detection of neurotransmitter-specific ^13^C labelling in mice and feasibility in the human brain at ultra-high field. POPE^13^C-MRS provides a readout of glutamatergic and GABAergic metabolism, enabling in vivo assessment of excitatory-inhibitory metabolic balance. This establishes a means for probing neurometabolic coupling with potential applications in translational and clinical neuroscience.

## Introduction

Neuroimaging techniques are essential to identifying metabolic biomarkers of brain function and disease, paving the way for personalised approaches to neuropsychiatric and neurodegenerative disorders^1^. While fluorodeoxyglucose positron emission tomography (FDG-PET) remains the gold standard for clinical assessment of cerebral glucose metabolism^2,3^, its focus on glucose uptake provides limited insight into downstream metabolic pathways that directly support neuronal signalling, including the glutamate-GABA cycle that underlies excitatory-inhibitory balance. Here, we introduce proton-observed proton-edited carbon-13 magnetic resonance spectroscopy (POPE^13^C-MRS), the first clinically compatible method enabling simultaneous indirect detection of glutamatergic and GABAergic ^13^C-labelling using standard ^1^H hardware. POPE^13^C-MRS uses targeted proton editing on standard proton (^1^H) MRI hardware to indirectly measure neurotransmitter-specific glucose metabolism by detecting ^1^H-^13^C satellite resonances, thereby enabling ^13^C labelling readout without broadband ^13^C excitation or the associated radiofrequency (RF) power deposition.

X-nuclear Magnetic Resonance Spectroscopy, which uses MRI-visible isotopes such as carbon-13 (^13^C) or deuterium (^2^H) to track labelled metabolic substrates, offers a powerful means to study neurometabolic pathways in vivo^4,5^. In particular, ^13^C-MRS following administration of ^13^C-labelled glucose allows real-time measurement of neurotransmission and energy metabolism via the tracking of label incorporation into the downstream metabolites^6,7^. However, despite this unique metabolic specificity, the implementation of ^13^C-MRS in human studies has been limited by three major technical challenges^8–12:^ its inherently low sensitivity (due to the lower gyromagnetic ratio of ^13^C compared to ^1^H), the requirement for high-power broadband radiofrequency (RF) excitation, and the need for specialised and costly RF hardware^13^.

Several strategies have been developed to address these limitations. Indirect ^13^C-MRS, such as proton-observed carbon-edited (POCE) ^13^C-MRS, improve sensitivity by detecting the protons attached to ^13^C-labeled metabolites, utilising the higher intrinsic sensitivity of protons^9,14,15^. However, these approaches rely on broadband ^13^C editing and decoupling, which substantially increases RF power deposition and SAR limiting their feasibility in human studies. Hyperpolarisation techniques provide dramatic gains in sensitivity by increasing nuclear spin polarisation far beyond thermal equilibrium levels^16,17^, enabling real-time metabolic imaging with unprecedented signal enhancement^18^. Yet this approach requires specialised equipment and infrastructure and is restricted to observing rapid metabolic processes due to the short (tens of seconds) lifetime of the hyperpolarised signal. Deuterium MRS (^2^H-MRS) has recently emerged as another promising strategy, enabling dynamic metabolic imaging (DMI) by exploiting the short longitudinal relaxation time of deuterium nuclei, allowing for rapid data sampling and higher signal-to-noise ratio^19^. However, it requires dedicated hardware and cannot resolve key neurotransmitter signals, as it does not distinguish glutamate from glutamine or detect GABA, metabolites that play central roles in excitatory and inhibitory neurotransmission through the glutamate-glutamine cycle between neurons and astrocytes^20,21^. More recently, indirect ‘dynamic’ labelling methods^22,23^, such as quantitative exchanged-label turnover (QUELT-MRS)^24^, have been proposed. These infer metabolite labelling from the decay of ^1^H resonances, when replaced by ^2^H or ^13^C, but rely on baseline scans and assumptions of steady-state metabolite pools following the labelled substrate administration, conditions that may not always hold during metabolic perturbations, such as those likely present in clinical populations.

Despite these advances, a straightforward and clinically deployable method for detecting ^13^C-labelling of glutamate and GABA in the human brain has remained elusive. Here, we develop and validate POPE^13^C-MRS, a method that tracks the incorporation of ^13^C from a labelled substrate (e.g. uniformly labelled [U-^13^C_6_]-glucose, which contains 6 ^13^C atoms) into downstream metabolites via measurable changes in proton MRS spectra, enabling non-invasive probing of excitatory, inhibitory, and glycolytic pathways in the brain (Fig.1). POPE^13^C-MRS uses a proton-edited sequence which is tailored to detect ^1^H signals from lactate H3 (LacH3), glutamate H4 (GluH4), GABA H4 and glutamate+glutamine H2 (GlxH2). Signals from these protons are only visible when they are bound to ^12^C, not ^13^C, meaning that these signals decrease as protons initially bound to ^12^C become coupled to ^13^C, as the [U-^13^C_6_]-glucose is metabolised, with the transferred signal appearing as characteristic ^13^C satellite resonances for selected metabolites.

**Figure 1:**
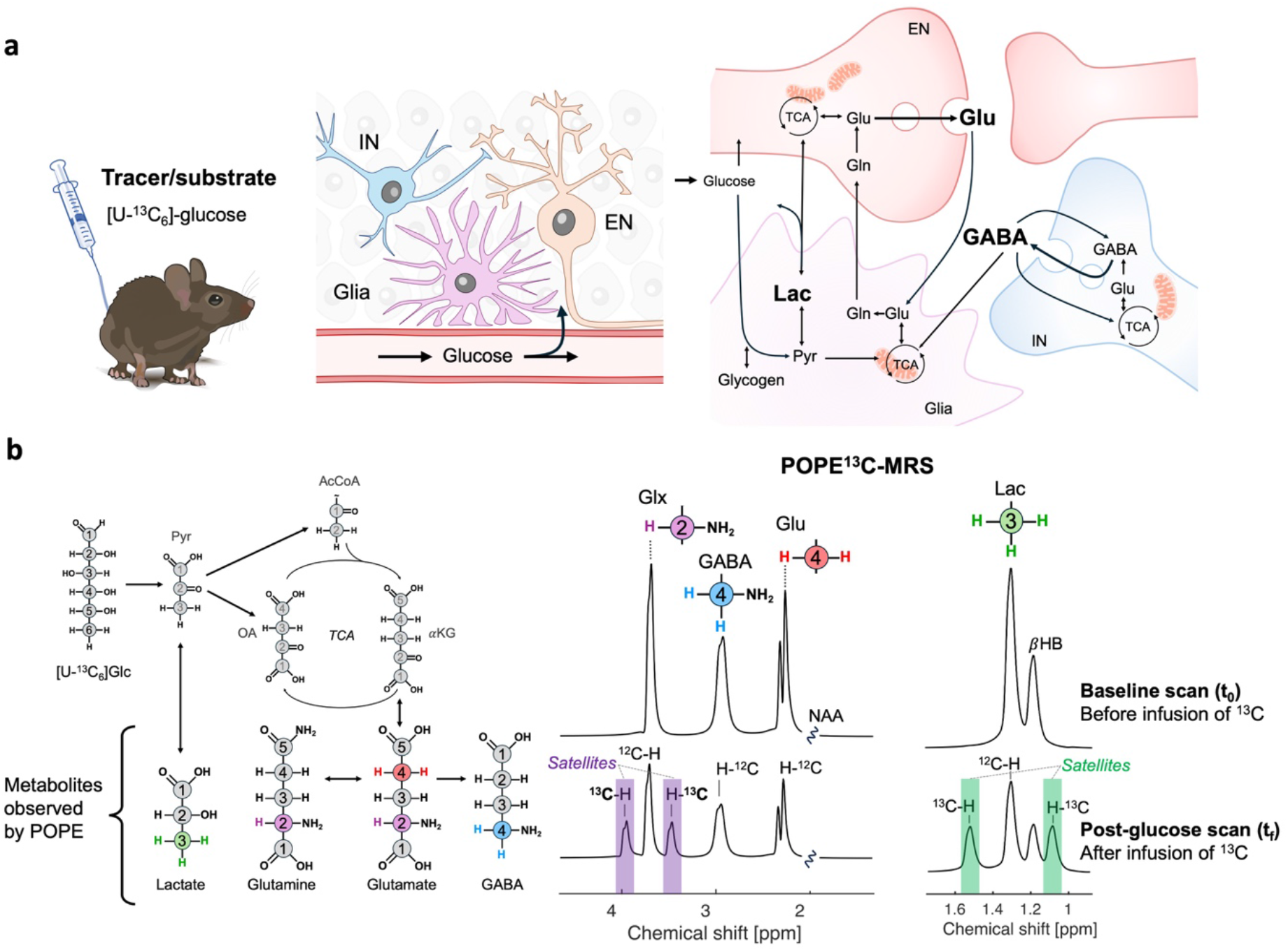
Theoretical background and metabolic basis of POPE^13^C-MRS. **a** Following administration of uniformly labelled ^13^C-glucose ([U-^13^C₆]Glc), ^13^C label is incorporated into lactate, glutamate, glutamine, and GABA through glycolysis, mitochondrial metabolism, and neurotransmitter cycling. Exchange between metabolic pools and compartments shapes the time course of ^13^C labelling. **b** POPE^13^C-MRS detects this incorporation indirectly via changes in ^1^H signals bound to ^12^C and the appearance of ^1^H-^13^C satellite resonances, enabling measurement of neurotransmitter-specific labelling. This approach provides a targeted readout of excitatory (glutamate) and inhibitory (GABA) metabolic pathway in vivo. [U-^13^C₆]Glc labels lactate via exchange with pyruvate generated by glycolysis, while glutamate and glutamine are labelled through exchange with tricarboxylic acid (TCA) cycle intermediates. GABA becomes labelled through the decarboxylation of glutamate. POPE^13^C-MRS detects protons (¹H) bound to ^12^C (or ^13^C) in specific positions in these metabolites, including glutamate and glutamine H2 (GlxH2, purple), glutamate H4 (GluH4, red), GABA H4 (GABAH4, blue), and lactate H3 (LacH3, green). As protons become bound to ^13^C, their ^1^H signal therefore decreases, producing a measurable signal loss. For some resonances (e.g., GlxH2 and LacH3 here), this loss is accompanied by the appearance of characteristic ^13^C-coupled satellite doublets flanking the parent resonance (see purple and green coloured boxes). Due to their lower signal and for the sake of simplicity, satellites of GABAH4 and GluH4 are not represented here. Carbon numbers in grey are not detectable with MRS due to the small metabolite pool size. IN, inhibitory neuron; EN, excitatory neuron; TCA, tricarboxylic acid cycle; Glc, glucose, Glu, glutamate; Gln, glutamine; Lac, lactate; Pyr, pyruvate; OA, oxaloacetate; aKG, alpha-ketoglutarate; AcCoA, acetyl-coenzyme-A; NAA, N-acetyl-aspartate; bHB, beta-hydroxybutyrate

We first establish and validate the method in pre-clinical models and then demonstrate its feasibility in the human brain. By enabling simultaneous and targeted measurement of excitatory and inhibitory metabolic pathways, POPE^13^C-MRS provides a method for investigating neurometabolic coupling in vivo, bridging a key gap between glucose uptake measurements and neurotransmitter-specific metabolism.

## Results

We established the theoretical foundation and optimised POPE^13^C-MRS in mice (Study 1), validated neurotransmitter labelling detection (Study 2), and assessed feasibility in the human brain (Study 3).

### Study 1: POPE^13^C-MRS general optimization

To test whether indirect detection of ^13^C-labelling with sensitivity to GABA and lactate could be achieved using proton-edited MRS, we administered [U-^13^C_6_]-glucose in mice while continuously acquiring MEGA-sLASER spectra. The results clearly demonstrate POPE^13^C-MRS can reliably detect the progressive incorporation of ^13^C into neurotransmitter pools through consistent changes in ^1^H-^12^C signal attenuation and emergence of ^1^H-^13^C satellite resonances (Fig.2). Because only a single echo time (TE) can be used for each edited sequence, we needed to optimize our TE for a single labelled resonance per spectrum. In the GABA-edited experiment, we selected GlxH2 as the optimization target because its ^13^C satellites were (i) most clearly resolved, (ii) minimally confounded by spectral overlap, and (iii) suitable as an internal reference for normalizing metabolite labelling. We then empirically determined the corresponding heteronuclear J-coupling constants (J_CH,_ the distance between the two satellite peaks) by sampling TE values around the expected theoretical optimum (with TE=N.1/2J, with N the number of J-revolutions). The J_CH_ constants are given by the distance between the two resonances of the doublet (in Hz), which was measured using a range of TEs, starting around established optimal values for GABA^25^ (68ms) and lactate (100ms, tested in phantom) (Supplementary Fig.2). We determined that a TE=69.5 ms (J_CH_(GlxH2)=143.9±2.7 Hz) for GABA-edited and TE=106 ms (J_CH_(LacH3)=122.7±1.0 Hz) for lactate-edited POPE^13^C-MRS should maximize the satellite detection. The J of GlxH2 was found to be slightly more variable, most likely because it is a combination of both J_CH_(GluH2) and J_CH_(GlnH2), which may not be identical, thus adding variability. Notably, the agreement between signal attenuation (GlxH2 and LacH3) and satellite appearance supports the internal consistency of the POPE^13^C-MRS readout (Fig.2c).

**Figure 2:**
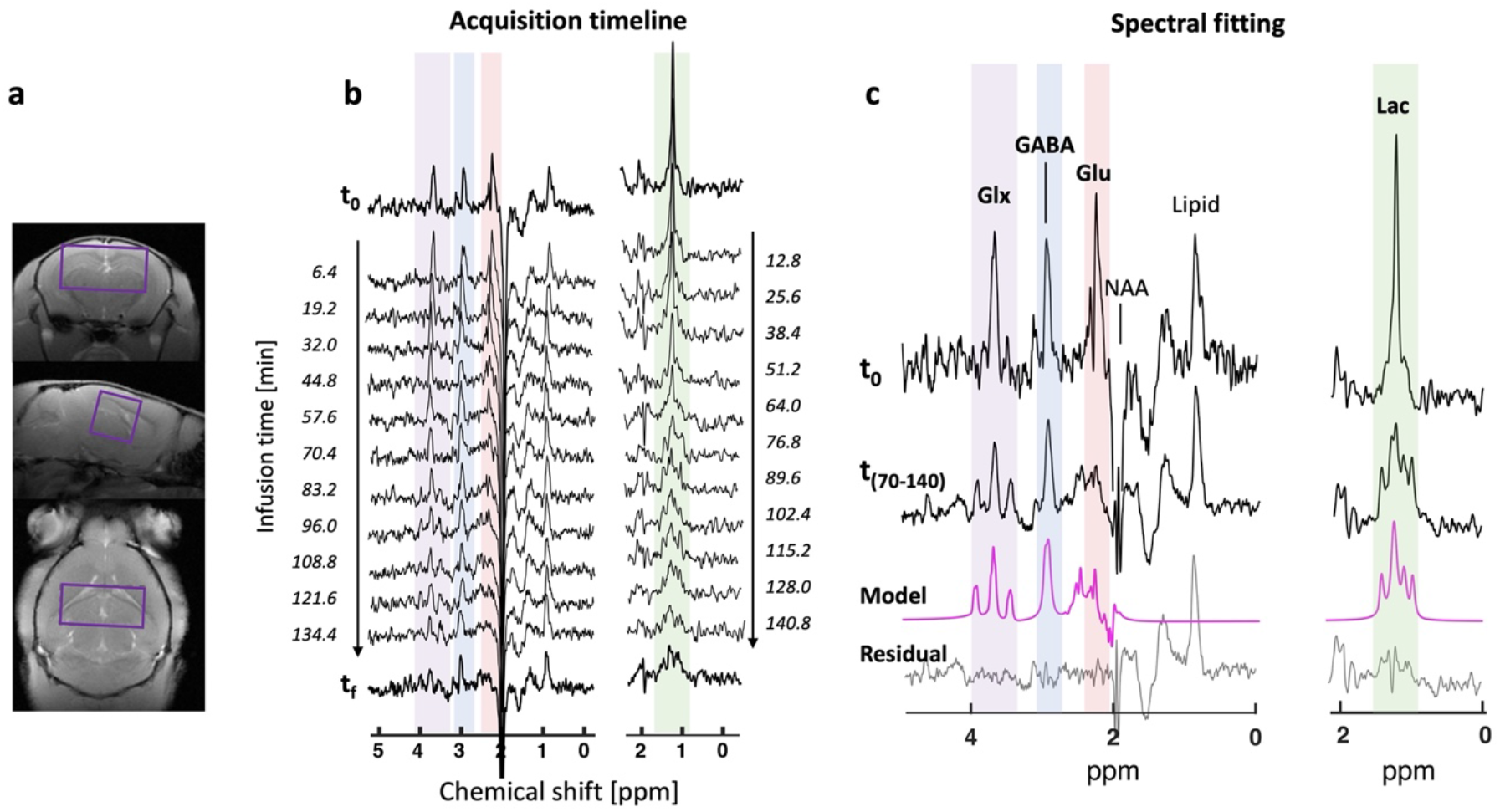
Time-resolved detection of neurotransmitter labelling following ^13^C-glucose administration. **a** Acquisition of POPE^13^C-MRS in mouse brain (in purple, voxel size: 6.5 x 3.4 x 3.2 mm^3^) following a ∼2h30 min s.c. administration of uniformly ^13^C-labelled glucose ([U-^13^C_6_]-glucose). **b** Progressive attenuation of ^1^H-^12^C resonances reflects incorporation of ^13^C label into downstream metabolites, consistent with metabolic flux through glutamatergic and GABAergic pathways. GABA- and lactate-POPE^13^C-MRS acquired from baseline scan (t_0_ = start of the infusion) until the end of the experiment (t_f_ = 2h30). **c** Fitting of POPE^13^C-MRS signals tracks neurotransmitter-specific metabolic labelling in vivo (averaged spectra acquired between 70- and 140-min post infusion). Peaks of interest include glutamate+glutamine-H2 (Glx) in purple and glutamate-H4 (Glu) in red, gamma-aminobutyric acid-H4 (GABA) in blue and lactate-H3 (Lac) in green. Spectra are all shown with a 5Hz Lorentzian apodization.

### Study 2: POPE^13^C-MRS detects ^13^C labelling of GABA, glutamate, glutamine and lactate after^13^C-glucose administration in mice

We then tested this protocol in Study 2 to assess the detectability and quantifiability of the satellites after a one-hour ^13^C-glucose infusion. Our data indicated that the ^13^C satellites of GlxH2 (^13^C-GlxH2) was consistently detectable across animals (Fig.3a) and could be quantified despite low labelling concentration (CRLB(GlxH2_tf_)=76%±27%, mean±s.d.; supplementary Fig.3). Measuring satellite signals are useful to control for change of total metabolite concentration throughout the infusion, a process that cannot be assessed by solely inferring the metabolite labelling from the drop of the initial ^1^H-^12^C signal. This was particularly relevant for Glx and lactate as we observed a 22±14% increase in total Glx and 13±5% in total lactate concentrations following one-hour infusion of ^13^C-Glc compared to the baseline scan in mice. Notably, the fitting of GluH4 and GlnH4 led to some residual signal likely due to GABAH2 resonance that could not be fitted and was thus not included in the basis set. Nonetheless, their CRLB values were in an acceptable range (GluH4_t0_ = 9%±4% and GlnH4_t0_ = 24%±33%). Interestingly, no lactate ^13^C-labelling was detected during the one-hour timeframe of the infusion, which was likely due to the use of medetomidine in these experiments, known to prevent lactate build up and labelling^26^ compared to isoflurane^27^, which was used in Study 1 (Fig.3b). While high lactate concentration and lactate ^13^C-satellites can be seen when using isoflurane, the use of medetomidine led to a complete absence of ^13^C-incorporation in the lactate pool (Supplementary Fig.4).

**Figure 3:**
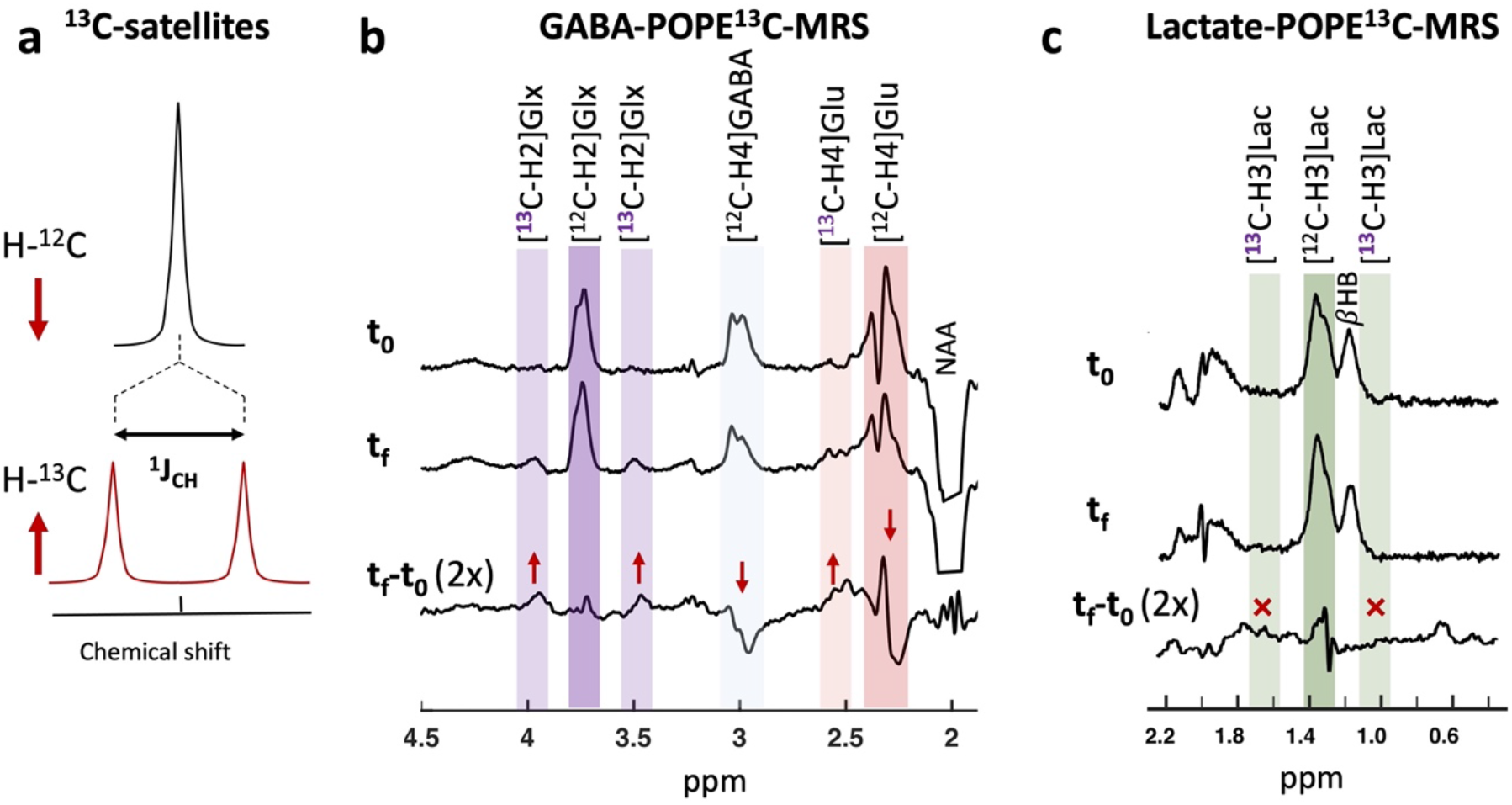
Validation of indirect detection of ^13^C-labelled metabolites. **a** Both attenuation of ^1^H-^12^C resonances and emergence of ^1^H-^13^C satellite signals provide complementary and internally consistent measures of metabolite labelling. **b** GABA-POPE^13^C-MRS can indirectly detect ^13^C labelling of GluC4 (via a drop in the ^12^C-GluH4 signal), GABAC4 (via a drop in the^12^C-GABAH4 signal) and of the total pool of glutamate and glutamine (via both the drop in ^12^C-GlxH2 signal and increase in satellite ^13^C-GlxH2 signals). Red arrows indicate the directionality of that change. **c** Lactate-POPE^13^C-MRS can detect indirect ^13^C labelling of lactate-C3 via both the drop in ^12^C-LacH3 signal and increase in satellite ^13^C-LacH3 signals (not detected here, red crosses). Experiments were performed in mice under medetomidine anaesthesia, making ^13^C-LacH3 signals undetectable. Spectra are averages of 5 adult mice after 1h [U-^13^C_6_]-glucose infusion and shown with 1 Hz Lorentzian apodization.

**Figure 4:**
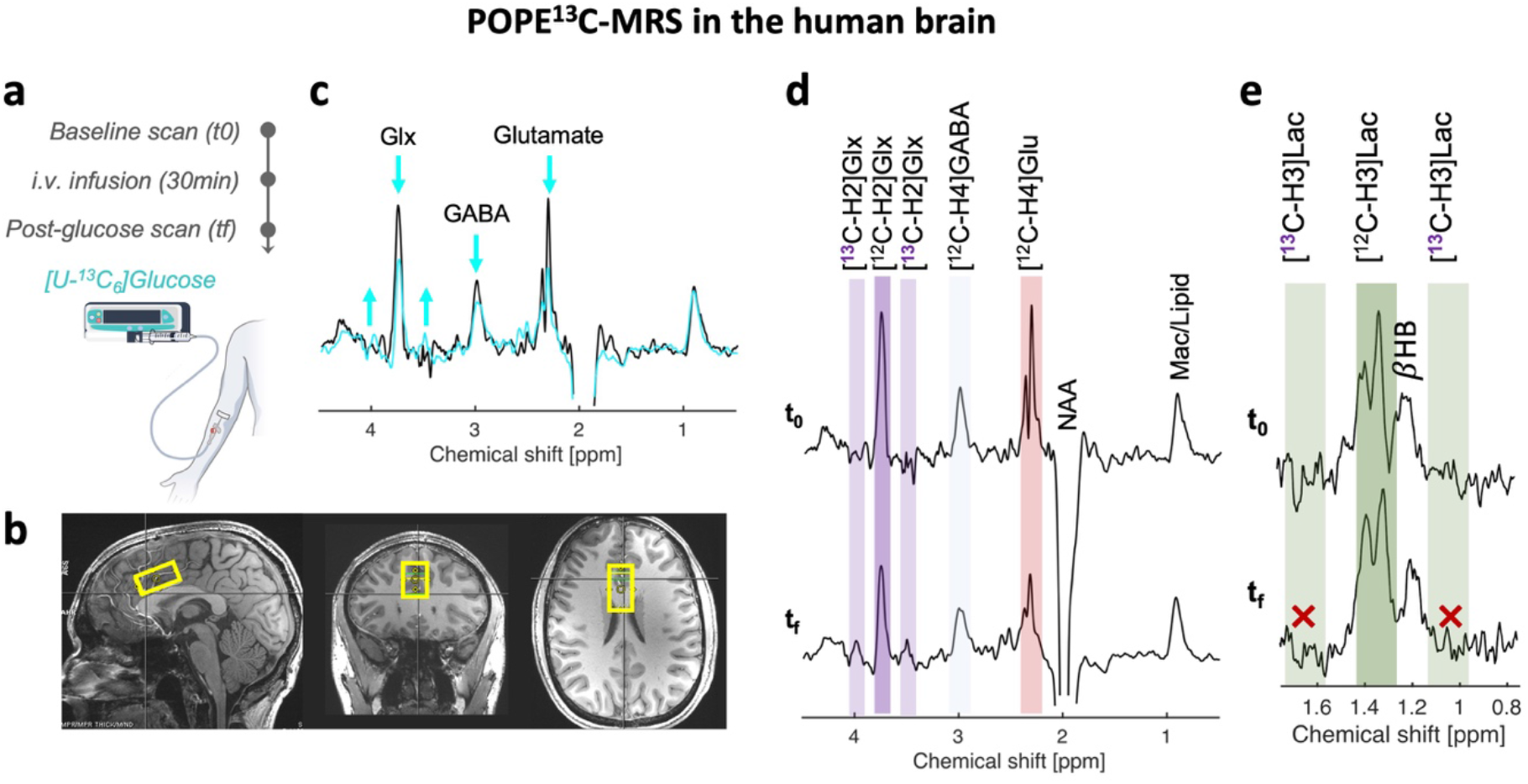
Feasibility of POPE^13^C-MRS in the human brain. **a** Timeline of the intravenous [U-^13^C_6_]-glucose infusion and POPE^13^C-MRS baseline (t0) and post-infusion (tf) acquisitions. **b** Voxel positioning in dorsal anterior cingulate cortex (dACC) for POPE^13^C-MRS acquisition. **c** Overlaid POPE^13^C-MRS before (black) and after (blue) the [U-^13^C_6_]-glucose infusion. **d,e** Detectable ^13^C-labelled GlxH2 satellite resonances and reduction in ^1^H-^12^C signals for GABA, Glu and Glx were observed following ^13^C-glucose administration, consistent with incorporation into neurotransmitter metabolic pools. Average GABA-POPE^13^C-MRS (d) and lactate-POPE^13^C-MRS (e) spectra before (baseline scan, t0) and after (post infusion scan, tf) the [U-^13^C_6_]-glucose administration. Glx = glutamate + glutamine; NAA, N-acetyl-aspartate; βHB, hydroxybutyrate; Mac/Lipid, macromolecules or lipids.

Within one hour of s.c. ^13^C-glucose infusion, the isotopic fractional enrichment (FE) of GlxH2 reached 0.21±0.06, while it was 0.36±0.09 for GABAH4 and 0.13±0.05 for GluH4. The comparatively high FE of GABA may reflect a combination of factors, including the relative metabolite pool sizes (Table 1; uncorrected concentrations: GABA: 4.40±1.07 [mM] vs. Glu: 3.99±0.71 [mM]) or differences in GABA metabolic turnover rates within the brain structures covered by the voxel (hippocampus and striatum). The amount of labelling of each metabolite pool is dependent on the ^13^C-glucose infusion time, thus metabolite labelling may be more comparable across groups or across experiments when reported relative to each other, such as by using the ^13^C-GluH4/^13^C-GABAH4 ratio. Alternatively, as the FE of GlxH2 (Glx_FE_) can be assessed with reference to its ^13^C-satellites, it can be used as a normalizing factor to report labelling of glutamate or GABA. The ^13^C-GluH4/^13^C-GABAH4 ratio here was thus 0.37±0.19, while the ^13^C-GluH4/Glx_FE_ was 2.82±1.42 and the ^13^C-GABAH4/Glx_FE_ was 8.32±3.77 (table 1). While coefficients of variation (CV) between animals were between ∼0.1-0.2 for basal metabolite concentrations, the CV were higher for the ^13^C-labelling concentrations (0.3-0.4), which is likely due to the variability in infusion efficiency across animals. Notably, using ^13^C-labelling ratios did not reduce inter-animal variability, consistent with the strong dependence of ratio stability on the arterial input function (AIF) and labelling dynamics, which we investigated below. Overall, these results confirm the feasibility of detecting neurotransmitter ^13^C-labelling within one hour of infusion in mice, while highlighting the critical influence of tracer delivery and labelling dynamics on quantification variability.

**Table 1:** Consistent detection of POPE^13^C-MRS readouts across mice. Quantification results from POPE^13^C-MRS in adult male mice (N=5) from Study 2. Metabolite quantifications are reported for baseline scans (t0) or after the [U-^13^C_6_]-glucose infusion (tf) using either water reference (given in millimolar, mM) or using the isotopic fractional enrichment of GlxH2 (Glx_FE_) as internal reference (given as arbitrary units, a.u.). Coefficient of variation (CV) represent the inter-individual variability of the quantification.

|  | GABA |  |  | Glu |  |  | Glu/GABA |  |
| --- | --- | --- | --- | --- | --- | --- | --- | --- |
| Time | $t_0$ | $t_f$ | | $t_0$ | $t_f$ | | $t_0$ | $t_f$ |
| Metabolite | [GABA] [mM] | $^{13}\text{C}$-GABA [mM] | $^{13}\text{C}$-GABA / Glx<sub>FE</sub> [a.u.] | [Glu] [mM] | $^{13}\text{C}$-Glu [mM] | $^{13}\text{C}$-Glu / Glx<sub>FE</sub> [a.u.] | [Glu/GABA] | [$^{13}\text{C}$-Glu/$^{13}\text{C}$-GABA] |
| Quantification | $4.40 \pm 1.07$ | $1.61 \pm 0.53$ | $8.32 \pm 3.77$ | $3.99 \pm 0.71$ | $0.52 \pm 0.21$ | $2.82 \pm 1.42$ | $0.92 \pm 0.08$ | $0.37 \pm 0.19$ |
| Coefficient of variation (CV) | 0.24 | 0.33 | 0.45 | 0.18 | 0.40 | 0.50 | 0.08 | 0.52 |

### Study 3: POPE^13^C-MRS detects brain neurotransmitter labelling in human brain

To evaluate the feasibility of POPE^13^C-MRS in humans, we tested the acquisition protocol in two participants to determine whether ^13^C labelling of brain GABA, glutamate, glutamine and lactate could be detected. Participants either received ^12^C-glucose (unlabelled) or ^13^C-glucose (labelled) intravenously. Each session included a baseline POPE^13^C-MRS scan in the dorsal anterior cingulate cortex (dACC), followed by the administration of 0.23 g/kg glucose over 30 minutes, and a post-infusion scan 1 hour later to allow for metabolic incorporation of ¹³C (Fig5.a,b). Following ^13^C-glucose administration, POPE^13^C-MRS detected reproducible ^13^Clabelled GlxH2 satellites resonances across acquisitions (Fig.5,c,d). The J-coupling constant of GlxH2 was measured and found to be similar to that of the mouse (Humans: J_CH_(GlxH2)≈146 Hz; Mouse: 144Hz). Quantification of the spectra indicated a 21% decrease in GABAH4 and 37% decrease in GluH4, after the ^13^C-glucose infusion (Table 2). Notably, GlxH2 signal dropped by 33%, but the GlxH2-satellites signals only grew by 19%. This discrepancy suggests suboptimal sensitivity of the current acquisition to the GlxH2 satellites in humans, likely reflecting a combination of imperfect TE optimization for human J-coupling values, low signal-to-noise (SNR), underestimated editing pulse profile, or physiological motion. Interestingly, no lactate ^13^C-labelling was observed (Fig.5e), probably due to the low conversion from glucose to lactate in human brain. Overall, these changes reflect incorporation of glucose-derived carbon into excitatory and inhibitory neurotransmitter pools, demonstrating the ability of POPE^13^C-MRS to probe neurotransmitter-specific metabolism in vivo.

**Figure 5:**
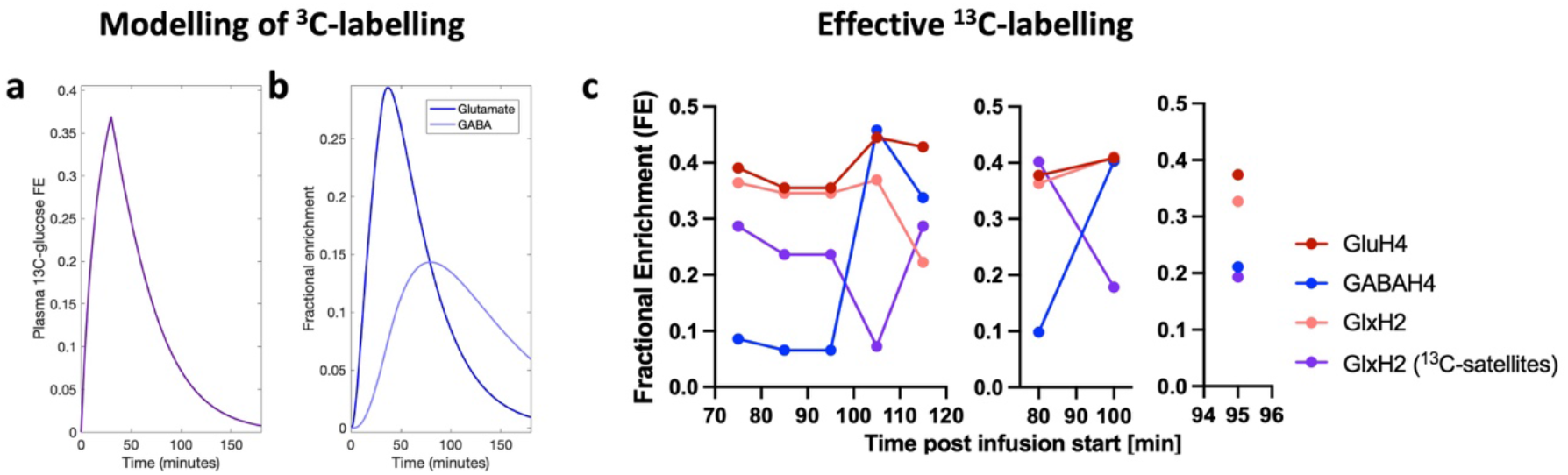
Assessment of metabolite labelling dynamics and ratio stability in human a,b. Simulation of arterial input function (AIF) based on our ^13^C-glucose infusion protocol (g) and estimation of the resulting ^13^C fractional enrichment (FE) of brain metabolites (h) in human POPE^13^C-MRS. **c** Effective ^13^C FE of metabolites measured with POPE^13^C in human dACC for different number of averages and SNR, i.e. (left) five individual acquisitions (SNR_average_=9.7), (middle) two averages (SNR_average_=9.8), and (right) a single average (SNR=11.6). GlxH2 FE was measured based on either the drop of basal GlxH2 peak from t_0_ (pink) or based on the GlxH2 satellite resonances (purple).

**Table 2:** POPE^13^C-MRS quantifications in the human. Quantification results from POPE^13^C-MRS in human in Study 3 (N=1). Metabolite quantifications are reported for baseline scans (t0) or after the [U-^13^C_6_]-glucose infusion (tf) using either water reference (given in millimolar, mM) or using the isotopic fractional enrichment of GlxH2 (Glx_FE_) as internal reference (given as arbitrary units, a.u.). Coefficient of variation (CV) represent the variability of the quantification within the ∼45min acquisition.

|  | GABA |  |  | Glu |  |  | Glu/GABA |  |
| --- | --- | --- | --- | --- | --- | --- | --- | --- |
| Time | $t_0$ | $t_f$ | | $t_0$ | $t_f$ | | $t_0$ | $t_f$ |
| Metabolite | [GABA] [mM] | $^{13}\text{C}$-GABA [mM] | $^{13}\text{C}$-GABA / Glx<sub>FE</sub> [a.u.] | [Glu] [mM] | $^{13}\text{C}$-Glu [mM] | $^{13}\text{C}$-Glu / Glx<sub>FE</sub> [a.u.] | [Glu/GABA] | $^{13}\text{C}$-Glu/$^{13}\text{C}$-GABA] |
| Quantification | 5.93 | 1.25 | 3.82 | 8.69 | 3.25 | 9.94 | 1.47 | 2.60 |
| Coefficient of variation (CV) |  | 0.90 | 0.96 |  | 0.10 | 0.30 |  | 0.63 |

The ^12^C-glucose session served as a control, allowing us to assess the stability and repeatability of POPE^13^C-MRS measurements in the absence of ^13^C labelling. By comparing the pre- and post-infusion scans, we could not identify changes in total metabolite levels due to sole administration of the 5% glucose bolus (Supplementary Fig.5), confirming that the drop in Glu and GABA observed with ^13^C-Glc are due to labelling and not changes in total metabolite pool concentration.

### Estimation of an optimal acquisition window for ^13^C-labelled metabolite ratios in POPE^13^C-MRS

Compared with time-resolved quantification^10,28^, metabolite labelling ratios offer a more practical approach for assessing excitatory–inhibitory metabolic balance. They improve SNR and allow for interruptions in acquisition – critical for the long scans required to resolve GABA labelling – but their stability, and hence comparability, is highly dependent on the ^13^C-labelling profile of the input tracer (supplementary Fig.6). We used a combination of experimental data and metabolic modelling to validate the stability and interpretability of metabolic labelling ratios accessible with POPE^13^C-MRS. Using ^13^C-MRS data in mouse brain under continuous ^13^C-glucose infusion, we sought to define the optimal acquisition window that maximizes the stability of metabolite ^13^C-labelling ratios and thereby minimizes sensitivity to variability in infusion protocols. We first evaluated ratio stability (Supplementary Fig.7) using theoretical labelling curves derived from a simplified pseudo–3-compartment model of brain metabolism (Supplementary Fig.7.a,d), compared to its associated original data (Supplementary Fig.7,e; data reused from previous work in Cherix et al.^28^), and then assessed temporal stability in the POPE^13^C-MRS data acquired here (Supplementary Fig.7).

Across all three ratios accessible with POPE^13^C-MRS (^13^C-GluH4/Glx_FE_, ^13^C-GABAH4/Glx_FE_ and ^13^C-GluH4/^13^C-GABAH4), stability increased with infusion duration and was maximal once labelling approached a plateau, whereas substantial variability was observed during the initial phase of the infusion (approximately the first hour). These findings highlight the importance of maintaining a stable, ideally near-linear arterial ^13^C-glucose input to promote constant metabolite ratios over time. Although theoretical modelling predicted the ^13^C-GluH4/^13^C-GABAH4 ratio to be the most stable across acquisition windows, this advantage was not fully realized in practice owing to the lower SNR of the GABA resonance, which introduced additional variability and could compromise glutamate labelling estimates. Together these results indicate that, when using continuous ^13^C-glucose infusion, labelling ratios are more reliable during later acquisition windows after labelling has stabilized, especially when GABA turnover is slow.

We next modelled the labelling profile expected from our human infusion protocol. The protocol comprised a 30-min bolus, a regime that is in theory suboptimal for maintaining stable metabolite labelling ratios (supplementary Fig.6e,f). Using the corresponding modelled arterial input function (Fig.5a), we predicted the resulting labelling dynamics of GluH4 and GABAH4 (Fig.5b) and compared these predictions with the empirically observed labelling profiles (Fig.5c). Notably, FE of Glu and Glx remained relatively stable across the acquisition window (80-100min post infusion), whereas GABA labelling increased substantially midway through the scan. This discrepancy between the model and the empirical data suggest that post-infusion glucose clearance and brain tracer availability may be slower or more prolonged in vivo than assumed by our model, likely reflecting physiological complexity not captured by the simplified AIF. Collectively, these findings indicate that, under our current human protocol, metabolite labelling curves approach a quasi-steady state approximately 2h after infusion onset, at which point labelling ratios may become reliable.

## Discussion

We introduce and validate POPE^13^C-MRS, a neuroimaging approach that enables simultaneous detection of glutamatergic and GABAergic metabolic labelling using standard MRI hardware. By combining molecular specificity with clinical compatibility, POPE^13^C-MRS addresses key practical constraints associated with indirect detection of neurotransmitter-specific ^13^C labelling in vivo. Using a cross-species validation approach, we demonstrate consistent detection and quantification of neurotransmitter-specific ^13^C labelling across mouse and human brain. This provides access to excitatory-inhibitory metabolic balance in vivo, a key feature of brain function that has been difficult to assess non-invasively. While in vivo GABA labelling in the human brain has been reported previously^10,29–31^, this is, to our knowledge, the first demonstration of detectable GABAergic labelling using an indirect ^13^C-MRS approach compatible with standard MRI hardware. The key advance lies in enabling simultaneous and internally consistent detection of excitatory and inhibitory metabolic labelling using a simple and widely accessible protocol. By enabling concurrent measurement of glutamatergic and GABAergic labelling, POPE^13^C-MRS provides direct access to excitatory-inhibitory metabolic balance, a central feature of brain function that has previously been difficult to assess non-invasively in humans.

Current clinical assessments of cerebral metabolism rely primarily on glucose-analogue PET imaging, which provides sensitive and quantitative measures of glucose uptake but offers no insight into downstream neurotransmitter-related metabolic pathways, including glutamate-glutamine cycling and GABA synthesis, which are altered in several neurological and psychiatric disorders. Emerging molecular MRI approaches, including hyperpolarized ^13^C methods^32^ and deuterium metabolic imaging^33^, allow pathway-specific interrogation of metabolic fluxes, yet their clinical adoption remains constrained by their requirement for specialized hardware and acquisition sequences or by their restricted metabolic coverage. Dynamic indirect ^13^C - and ^2^H-MRS offer a complementary strategy to probe downstream neuroenergetic and neurotransmitter-related processes while remaining compatible with standard MRI platforms^22–24,31^. POPE^13^C-MRS builds on this line of methodological developments but is specifically tailored to enable reliable access to GABAergic metabolism, which has remained technically challenging with existing indirect approaches. By refining widely available ¹H-MRS techniques and leveraging standard radiofrequency hardware, POPE^13^C-MRS addresses key limitations of direct detection at ^2^H or ^13^C frequencies, which typically require dedicated coils and tailored pulse sequences. The method is based on a standard proton editing sequence (MEGA-sLASER), making it readily deployable in clinical research environments. Similar to recent indirect ^13^C approaches (e.g. selPOCE) that prioritize targeted detection of selected resonances to infer neurotransmission cycling^9,34^, POPE^13^C-MRS deliberately trades breadth of metabolic coverage for robustness and specificity, focusing on metabolites most informative for excitatory–inhibitory metabolism. In this context, POPE^13^C-MRS is not intended as a general-purpose ^13^C-MRS replacement, but as a targeted tool optimized for probing GABAergic metabolism alongside glutamatergic and glycolytic pathways. Using this approach, we reliably detected labelling of GABAH4, GluH4 and GlxH2 in both mouse and human brain.

Isotopic labelling of GABA and lactate in the human brain has been reported previously^29–31;^ however, detection of GABA has been severely constrained by its slow metabolic turnover and strong spectral overlap with neighbouring resonances. As a result, reported FE of GABA in human brain have typically been low (<12%), often approaching the noise floor of conventional acquisition protocols. In contrast, POPE^13^C-MRS enabled reliable detection of GABAH4 labelling within 100 minutes after infusion onset, reaching a mean FE of ∼21% over the average acquisition window. The use of editing-based ^1^H-MRS not only improves measurement precision for GABA and lactate, but also enhances applicability at lower field strengths, where specific absorption rate constraints associated with broadband editing pulses can be less restrictive. This opens a pathway for translating POPE^13^C-MRS to more common 3 Tesla systems, potentially broadening applicability while preserving sufficient SNR for edited metabolite detection. While animal studies benefit from a broader range of ^13^C-MRS methodologies that offer higher metabolic coverage and are less constrained by SAR limitations, the simplicity and clinical compatibility of POPE^13^C-MRS may also facilitate translational pipelines by harmonizing protocols across species.

The ability to quantify GABA, glutamate and lactate labelling with improved precision provides biologically meaningful access to key components of brain metabolism. GABA is the principal inhibitory neurotransmitter in the mammalian brain, and imbalances between excitatory and inhibitory metabolic activity are implicated in a wide range of neurological and psychiatric disorders^35,36^. Importantly, GABA is particularly abundant in deep brain structures such as the thalamus or striatum^37–39^, which are difficult to probe with conventional MRS approaches. By enabling indirect ^13^C measurements in these regions, POPE^13^C-MRS may extend metabolic imaging beyond cortical targets that have dominated prior ^13^C-MRS work^10^. Lactate, as a major end-product of glycolysis, reflects aerobic glycolysis and astrocytic metabolic activity, processes that remain incompletely understood in the human brain^40^. In our study, however, ^13^C-labelling of lactate was not detectable in human, consistent with previous reports of low conversion of glucose into lactate under normal physiological conditions^11,41^. Notably, higher levels of lactate labelling from glucose have been reported in several pathologies^42–44^, suggesting that POPE^13^C-MRS may provide a sensitive tool to detect such metabolic alterations. Because lactate labelling depends not only on glucose uptake but also on astrocytic glycogen turnover^45^, which may causes an apparent dilution of ^13^C-lactate signal, the POPE^13^C-MRS may provide indirect sensitivity to astroglial metabolism and glycogen shunting, thereby offering new opportunities to study neurometabolic coupling in vivo. Consequently, POPE^13^C-MRS may be particularly well suited for clinical investigations of disorders characterised by dysregulated cerebral metabolism, including conditions associated with brain insulin resistance and altered excitatory–inhibitory balance.

Despite these advances, several technical challenges and limitations remain. While sensitivity and prolonged acquisitions required to resolve GABA labelling currently limit full metabolic fluxes modelling, the ability to derive stable metabolic labelling ratios provide a practical and biologically meaningful proxy for excitatory-inhibitory metabolic balance in vivo. However, such ratios are inherently dependent on the tracer administration protocol, arterial input function, and the temporal dynamics of metabolite labelling. Because labelling curves typically follow exponential saturation rather than linear trajectories, ratio stability can only be expected within specific temporal windows. We therefore identified acquisition windows in which labelling ratios are sufficiently stable to provide reliable and interpretable readouts, emphasizing the need for protocol harmonization to enable meaningful comparisons across studies. FE of lactate has previously been used as AIF in animal studies^46,47^; however, this was not feasible under light anaesthesia or in human herein, necessitating alternative normalization strategies to improve the comparability of labelling measures. We therefore corrected metabolite labelling by the FE of ^13^C-GlxH2, which was expected to be less sensitive to specific excitatory or inhibitory flux changes than ^13^C-GABAH4 or ^13^C-GluH4. Alternatively, the ^13^C-GluH4/^13^C-GABAH4 ratio provided a comparatively stable metric over prolonged infusion periods, reducing sensitivity to infusion timing. An additional source of variability may arise from the contamination of the GABAH4 signal by co-edited macromolecule (MM) resonances, such that the edited signal reflects GABA+ rather than GABA alone^48^. This macromolecular contribution can vary across subjects and physiological states and may therefore contribute to variability in apparent labelling across studies.

A further methodological consideration concerns the quantification of ^13^C satellite resonances, which are inherently challenging without appropriate calibration. This difficulty arises from the fast heteronuclear J-evolution (with J_CH_-coupling constants typically ranging from 130 to 160 Hz), which can cause signal dephasing, and from the reduced SNR due to the splitting of the original signal into two components. Many indirect ^13^C-MRS approaches rely on baseline scans acquired prior to tracer administration, implicitly assuming stable total metabolite pool sizes. While this assumption held in our human control experiments with unlabelled glucose, we observed changes in total Glx and lactate pools in mice following glucose administration, indicating that this assumption may not hold in all physiological or pathological contexts. Although ^13^C satellite detection can, in principle, provide access to changes in total metabolite pools, their reliable measurement requires careful calibration of echo times and editing pulse profiles. In humans, we observed a small discrepancy between GlxH2 labelling estimated from peak disappearance and from satellite resonances appearance, likely reflecting imperfect sequence calibration and subtle differences in effective J-coupling constants between species. Notably, the use of sLASER rather than PRESS likely proved critical for robust satellite detection, owing to improved refocusing homogeneity of fast-evolving satellite resonances with adiabatic pulses^49^.

Future work should focus on improving sensitivity, optimising tracer delivery, and validating reproducibility in larger human cohorts. These developments will be essential to establish POPE^13^C-MRS as a robust tool for investigating neurometabolic coupling and excitatory–inhibitory balance in clinical populations.

## Materials and Methods

POPE^13^C-MRS experiments were conducted in mice and humans to evaluate sequence optimisation, metabolite labelling detectability, and cross-species feasibility.

### Study 1 and 2 (mouse)

#### Animals

Male and Female C57BL6/J mice were used at the age of 3 months. Animals were housed in standard conditions (12h day-light cycle, 20-24°C, 46-65% humidity). Animals had *ad libitum* access to standard rodent chow diet and water. All procedures were approved by the University of Oxford ethical review committee and conducted under UK Home Office regulations (Animal [Scientific Procedures] Act 1986), and in compliance with the ARRIVE (Animal Research: Reporting *in vivo* Experiments) guidelines.

#### Anaesthesia and monitoring

Anaesthesia was induced with a bolus of 4% isoflurane mixed with air after which a single bolus of medetomidine in saline (0.1 mg/ml) was administered (0.2 mg/kg, s.c.) followed by a continuous infusion (0.4 mg/kg/h, s.c.) with minimal isoflurane administration (0.2% in air containing 35% O_2_). Some animals were only kept under 1-1.5% isoflurane following the bolus for the protocol optimization. The breathing rate and rectal temperature were monitored during the entire scans using a small animal monitoring system (SA Instruments Inc., New York, USA).

#### ^13^C-glucose Infusion

After the baseline POPE^13^C-MRS scan (see below), animals were administered a (20% w/v) solution of (99%) uniformly ^13^C-labelled glucose ([U-^13^C_6_]-Glc) solution (CK Isotopes Ltd.) via a sub-cutaneous (s.c.) cannula connected with an infusion pump (PHD 2000, Harvard Apparatus) which was inserted in the animal’s flank. A subcutaneous infusion was used to achieve slower absorption kinetics and approximate linear increase in blood glucose ^13^C-labelling, which consisted of a 5 min bolus (2.83 mL/kg, 99% FE) followed by continuous flow (10 mL/kg/h, 99% FE).

#### MRI/MRS acquisition

POPE^13^C-MRS was acquired on a 7T (70/20) BioSpec MRI scanner (Bruker, Ettlingen, DE) equipped with an 86 mm transmit volume coil and 2×2 receive ^1^H-cryoprobe and using Paravision 360.1.1. Anatomical T_2_-weighted MRI images were acquired for the MRS voxel placement with a localizer (FLASH, TE/TR= 3ms/15ms, 2 averages). The MRS voxel (6.5 × 3.4 × 3.2 mm^3^) encompassed the dorsal hippocampal area and striatum at the centre-top of the brain. Shimming (MAPSHIM) was performed in the voxel to reach a water Full Width at Half Maximum (FWHM) below 15Hz. POPE^13^C-MRS data were acquired using an in-house implementation of a MEscher-GArwood semi-Localization by Adiabatic Selective Refocusing (MEGA-sLASER)^50^ sequence, configured to selectively detect GABA/glutamate or lactate. Editing pulse offsets and bandwidths were optimised for each metabolite (see Supplementary Methods).

#### Data processing and modelling

Spectra were processed using jMRUI and quantified with QUEST^51^ (see Supplementary Methods). ON and OFF transients were phase- and frequency-corrected and averaged separately before subtraction (ON-OFF). Baseline (t_0_) and post-infusion (t_f_) spectra were then aligned, apodised (2Hz), removal of the NAA signal using Hankel-Lanczos Singular Value Decomposition (HLSVD)^52^, and analysed using predefined basis set (Supplementary Fig.1). Absolute concentrations were estimated using the unsuppressed water signal as an internal reference (80% brain water content ^53^).

To assess the reliability and consistency of metabolite ^13^C-labelling ratios, we used raw data from Cherix et. al.^28^ acquired on a horizontal 14.1 T scanner in mouse brain using indirect ^13^C-MRS (^1^H[^13^C]-MRS) as previously described^54,55^. The labelling curves fitted from this dataset using a pseudo 3-compartment model was then used to estimate how the theoretical labelling ratios (e.g. ^13^C-GluC4/^13^C-GABAC4) fluctuate throughout this experiment under a continuous ^13^C-Glc infusion. A similar approach was then used for the data acquired herein with POPE^13^C-MRS in mice. To determine how ^13^C-labelling ratios are affected by the acquisition window, i.e. the timing between the start and end of the acquisition relative to the start of the ^13^C-Glc infusion protocol, the average ratio was computed for variable windows and was reported relative to its final ratio value.

#### Study 3 (human)

Experiments in humans conformed to the Declaration of Helsinki, apart from pre-registration, and were approved by the Medical Sciences Interdivisional Research Ethics Committee of the University of Oxford (MS IDREC-R94883/RE001). Two healthy male participants (27 and 33 years old) were recruited and provided written informed consent. Neither participant had a history of diabetes, hyperglycaemia, neurological or psychiatric conditions and both met local MRI safety criteria. Participants had a one-hour scan, followed by a ^13^C-glucose infusion and a second one-hour scan.

#### Infusion protocol

^13^C-labelled D-glucose ([U-^13^C_6_]Glc, 99% ^13^C enrichment) was purchased (CK Isotopes Limited) and processed for intravenous (i.v.) infusion in accordance with EU cGMP principles. Participants were asked to fast for 8 hours prior to the experiment. Participants received either ^13^C-glucose or unlabelled (^12^C) glucose (0.23 g/kg, 5% w/v)^2^ via intravenous infusion over 30 min. The remaining isotope was flushed through the infusion line using isotonic saline.

#### MRI/MRS acquisition

Participants were scanned in a Siemens MAGNETOM 7T system (Siemens Healthineers, Erlangen, Germany) with a Nova Medical 8Tx32Rx head coil (Nova Medical Inc, Wilmington, MA, USA). An anatomical T1w MPRAGE image was acquired for placing the MRS voxel (TR=2.6 s, TE=3.31 ms, isotropic resolution 0.7 x 0.7 x 0.7 mm, 256 slices). The MRS voxel was positioned in the dorsal anterior cingulate cortex (dACC, 16 x 16 x 35 mm). First- and second-order shims were adjusted first by vendor-implemented gradient-echo shimming (“GRE Brain”)^56^, then fine adjustment of shims were done using FAST(EST)MAP^57^. A MEGA-sLASER sequence (CMRR Spectro Package, University of Minnesota^58^) was used (TR = 4s, 2048 points, acquisition BW = 3kHz, number of transients =32 x 2 editing conditions, editing bandwidth = 160Hz) for GABA (TE = 69.5 ms, editing offset 1.90 ppm and 7.50 ppm) and lactate (TE = 106 ms, editing offset 4.10 ppm and 5.10 ppm). Water suppression was achieved with VAPOR^59^ (150Hz Bandwidth) and outer volume saturation (OVS) was enabled on the anterior-posterior axis. The MRS acquisition consisted of five GABA-POPE^13^C-MRS and five Lactate-POPE^13^C-MRS segments acquired in interleaved sequence, with an unsuppressed water reference signal acquired at the beginning and the end of the MRS protocol. An identical protocol was used at baseline (t0; before the ^13^C-Glc infusion) and final (tf; after the ^13^C-Glc infusion) scans.

#### Data processing and modelling

MRS data was processed as described above for the mouse data, i.e. using QUEST routine^60^ in jMRUIv6.0^51^ using a combination of basis sets with identical prior knowledge.

Plasma [U-¹³C₆]glucose enrichment following a 30 min constant intravenous infusion of 5% glucose (99% ¹³C; 3.2 mmol min⁻¹) in our participant was modelled using a finite-duration input function with first-order clearance, 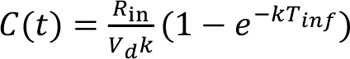 during infusion and mono-exponential washout thereafter 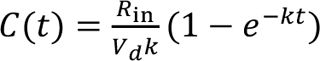. The apparent distribution volume of glucose was estimated^61^ to be *V_d_* = 0.26 L kg^-1^ and a first-order fractional clearance rate of plasma glucose was estimated^62^ to be *k* = 0.029 min^-1^. Plasma fractional enrichment *AIF*(*t*) was computed assuming clamped total glucose (baseline 5 mmol L⁻¹) and 99% tracer purity. Brain metabolite labelling was modelled using first-order driven kinetics to estimate the labelling dynamics of GABAH4 and GluH4 within a simplified representation of cerebral metabolism (Equations 1 and 2). Equation 1 describes glutamate labelling dynamics, and Equation 2 describes GABA labelling dynamics, where effective influx and efflux constants represent composite metabolic fluxes derived from literature values.

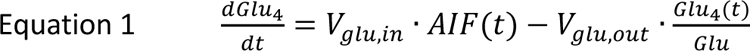

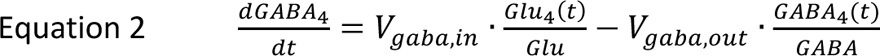

Effective influx and efflux constants *V*_Glu,in_ = 0.86 µ*mol g*^-1^*min*^-1^, *V*_Glu,out_ = 0.95 µ*mol g*^-1^*min*^-1^and *V*_G*ABA*,*in*_ = *V*_G*ABA*,*o*ut_ = 0.086 µ*mol g*^-1^*min*^-1^ were defined as composite fluxes including literature-based values for human brain. Glutamatergic influx *V*_g*lu*,*in*_ was defined as the sum of transmitochondrial (*V_x_*) and excitatory neurotransmission (*V_NT_^e^*) fluxes, while glutamatergic efflux also included glutamate decarboxylation (*V_GAD_*), i.e. *V*_g*lu*,*in*_ = *V_x_* + *V_NT_^e^* and *V*_g*lu*,*out*_ = *V_x_* + *V_NT_^e^* + *V_GAD_*, with *V_x_* = 0.57 µ*mol g*^-1^*min*^-1^,^63^ *V_NT_^e^* accounting for 90% of total neurotransmission cycling^29,64^ *V_NT_* = 0.32 µ*mol g*^-1^*min*^-1^and *V_GAD_* accounting for 10% of *k*_G*lu*,*in*_ (*V_GAD_* = 0.086 µ*mol g*^-1^*min*^-1^).^15,29^ GABAergic influx was set equal to its efflux, with glutamate decarboxylation being equivalent to the sum of GABAergic neurotransmission (*V_NT_^i^* = 0.1 · *V_NT_* = 0.032 µ*mol g*^-1^*min*^-1^),) and GABA recycling (*V_s_*_ℎ*unt*_*^i^* = 0.054 µ*mol g*^-1^*min*^-1^), i.e. *V*_g*aba*,*in*_ = *V_GAD_* and *^V^*_g*aba*,*out*_ ^= (*V*^*_NT_^i^*^+*V*^*_s_*_ℎ*unt*_*^i^*^) = *V*^*_GAD_*.

## Statistics

Statistics were all performed with GraphPad Prism (GraphPad Software, San Diego, CA, USA). All values are given as mean± standard deviation. unless stated otherwise. The ^13^C-labelled metabolite ratio analysis and human ^13^C-labelling dynamics were performed using MATLAB (R2020a).

## Acknowledgements

This study was supported financially by the Swiss National Science Foundation (P500PM_203208 to AC) and the University of Oxford Medical Sciences Internal Fund (Pump-Priming, 0014920 to AC). The Wellcome Centre for Integrative Neuroimaging (WIN, now OxCIN) was supported by core funding from the Wellcome Trust (203139/Z/16/Z and 203139/A/16/Z). This work was supported by the NIHR Oxford Health Biomedical Research Centre (NIHR203316). The views expressed are those of the author(s) and not necessarily those of the NIHR or the Department of Health and Social Care. We also thank the UK BBSRC (grant number BB/W019582/1) for support. WTC is funded by Wellcome [225924/Z/22/Z]. CJS holds a Wellcome Trust Senior Research Fellowship [224430/Z/21/Z].

The MRS package was developed by Edward J. Auerbach and Małgorzata Marjańska and provided by the University of Minnesota under a C2P agreement. We thank Damian Tyler (OCMR Oxford) for advice and feedback on the manuscript. We thank Sean Smart (OxCIN core staff) for support with preclinical equipment. We thank Roswell Shelhamer (Cambridge Isotope Laboratories, Inc.) and Ben Pepper (CK isotopes Ltd.) for their helpful discussions and support in sourcing the isotope. We acknowledge Rolf Gruetter for constructive feedback on the manuscript and for supporting the reuse of previously acquired data.

This research was funded in whole, or in part, by the Wellcome Trust [Grant number 203139/Z/16/Z, 203139/A/16/Z, 224430/Z/21/Z, and 225924/Z/22/Z]. For the purpose of open access, the author has applied a CC BY public copyright licence to any Author Accepted Manuscript version arising from this submission.

## Authors’ contributions

AC designed the study. AC, MT, WC and JC optimized the protocol. AC, JC, MT and OH acquired the data. AC analysed and interpreted the data. AC drafted the manuscript. All the authors assisted in revising the manuscript and approved the final version.

## Ethics Declaration

The authors declare no conflicts of interest with respect to the research, authorship, and/or publication of this article.

## Data availability statement

Source data underlying the figures are provided as a Source Data file. Additional data supporting the findings of this study are available from the corresponding author upon reasonable request.

## References

1. Abi-Dargham, A. & Horga, G. The search for imaging biomarkers in psychiatric disorders. Nat. Med. 22, 1248–1255 (2016).

2. Patil, S. et al. Molecular Imaging with PET in the Assessment of Vascular Dementia and Cerebrovascular Disease. PET Clin. 20, 121–131 (2025).

3. Salmon, E., Collette, F. & Bastin, C. Cerebral glucose metabolism in Alzheimer’s disease. Cortex 179, 50–61 (2024).

4. Lizarbe, B., Cherix, A. & Gruetter, R. In vivo heteronuclear magnetic resonance spectroscopy. Methods Mol. Biol. 1718, 169–187 (2018).

5. Straathof, M., Meerwaldt, A. E., De Feyter, H. M., de Graaf, R. A. & Dijkhuizen, R. M. Deuterium Metabolic Imaging of the Healthy and Diseased Brain. Neuroscience 474, 94–99 (2021).

6. Gruetter, R. et al. Localized *in vivo*^13^ C NMR spectroscopy of the brain. NMR Biomed. 16, 313–338 (2003).

7. Rothman, D. L., De Feyter, H. M., De Graaf, R. A., Mason, G. F. & Behar, K. L. ^13^ C MRS studies of neuroenergetics and neurotransmitter cycling in humans. NMR Biomed. 24, 943–957 (2011).

8. Gruetter, R. et al. Localized 13C NMR Spectroscopy in the Human Brain of Amino Acid Labeling from [1-13C]Glucose. J. Neurochem. 63, 1377–1385 (1994).

9. De Feyter, H. M. et al. Selective proton-observed, carbon-edited (selPOCE) MRS method for measurement of glutamate and glutamine^13^ C-labeling in the human frontal cortex. Magn. Reson. Med. 80, 11–20 (2018).

10. Abdallah, C. G. et al. Glutamate metabolism in major depressive disorder. Am. J. Psychiatry 171, 1320–7 (2014).

11. Chhina, N. et al. Measurement of human tricarboxylic acid cycle rates during visual activation by^13^ C magnetic resonance spectroscopy. J. Neurosci. Res. 66, 737–746 (2001).

12. Moreno, A., Ross, B. D. & Blüml, S. Direct determination of the *N*-acetyl-L-aspartate synthesis rate in the human brain by^13^ C MRS and [1-^13^ C]glucose infusion. J. Neurochem. 77, 347–350 (2001).

13. de Graaf, R. A., Rothman, D. L. & Behar, K. L. State of the art direct 13C and indirect 1H-[13C] NMR spectroscopy in vivo. A practical guide. NMR Biomed. 24, 958–72 (2011).

14. Pfeuffer, J. et al. Localized in vivo 1H NMR detection of neurotransmitter labeling in rat brain during infusion of [1-13C] D-glucose. Magn. Reson. Med. 41, 1077–83 (1999).

15. van Eijsden, P., Behar, K. L., Mason, G. F., Braun, K. P. J. & de Graaf, R. A. In vivo neurochemical profiling of rat brain by 1H-[13C] NMR spectroscopy: cerebral energetics and glutamatergic/GABAergic neurotransmission. J. Neurochem. 112, 24–33 (2010).

16. Le Page, L. M., Guglielmetti, C., Taglang, C. & Chaumeil, M. M. Imaging Brain Metabolism Using Hyperpolarized 13C Magnetic Resonance Spectroscopy. Trends Neurosci. 43, 343–354 (2020).

17. Jørgensen, Sh., et al. Hyperpolarized MRI – An Update and Future Perspectives. Semin. Nucl. Med. 52, 374–381 (2022).

18. Grist, J. T. et al. Hyperpolarized^13^ C MRI: A novel approach for probing cerebral metabolism in health and neurological disease. J. Cereb. Blood Flow Metab. 40, 1137–1147 (2020).

19. De Feyter, H. M. et al. Deuterium metabolic imaging (DMI) for MRI-based 3D mapping of metabolism in vivo. Sci. Adv. 4, eaat7314 (2018).

20. Westergaard, N., Sonnewald, U. & Schousboe, A. Metabolic Trafficking between Neurons and Astrocytes: The Glutamate/Glutamine Cycle Revisited. Dev. Neurosci. 17, 203–211 (1995).

21. Rothman, D. L. et al. In vivo nuclear magnetic resonance spectroscopy studies of the relationship between the glutamate--glutamine neurotransmitter cycle and functional neuroenergetics. Philos. Trans. R. Soc. Lond. B. Biol. Sci. 354, 1165–1177 (1999).

22. Boumezbeur, F. et al. NMR measurement of brain oxidative metabolism in monkeys using13C-labeled glucose without a13C radiofrequency channel. Magn. Reson. Med. 52, 33–40 (2004).

23. Dehghani, M., Zhang, S., Kumaragamage, C., Rosa-Neto, P. & Near, J. Dynamic 1 H-MRS for detection of 13 C-labeled glucose metabolism in the human brain at 3T. Magn. Reson. Med. 84, 1140–1151 (2020).

24. Bednarik, P. et al. 2H labeling enables non-invasive 3D proton MR imaging of glucose and neurotransmitter metabolism at 7T in the human brain. *Nat*. Biomed. Eng. 7, 1001–1013 (2023).

25. Rothman, D. L., Petroff, O. A., Behar, K. L. & Mattson, R. H. Localized 1H NMR measurements of gamma-aminobutyric acid in human brain in vivo. Proc. Natl. Acad. Sci. 90, 5662–5666 (1993).

26. Flatt, E. et al. Measuring Glycolytic Activity with Hyperpolarized [2H7, U-13C6] D-Glucose in the Naive Mouse Brain under Different Anesthetic Conditions. Metabolites 11, 413 (2021).

27. Cherix, A. et al. Deletion of Crtc1 leads to hippocampal neuroenergetic impairments associated with depressive-like behavior. Mol. Psychiatry (2022) doi:10.1038/s41380-022-01791-5.

28. Cherix, A. et al. Excitatory/inhibitory neuronal metabolic balance in mouse hippocampus upon infusion of [U-13 C 6]glucose. J. Cereb. Blood Flow Metab. 632, 0271678X2091053 (2020).

29. Gruetter, R., Seaquist, E. R., Kim, S. & Ugurbil, K. Localized in vivo^13^C-NMR of Glutamate Metabolism in the Human Brain: Initial Results at 4 Tesla. Dev. Neurosci. 20, 380–388 (1998).

30. Bednarik, P. et al. 1H magnetic resonance spectroscopic imaging of deuterated glucose and of neurotransmitter metabolism at 7 T in the human brain. *Nat*. Biomed. Eng. (2023) doi:10.1038/s41551-023-01035-z.

31. Rich, L. J. et al. 1H magnetic resonance spectroscopy of 2H-to-1H exchange quantifies the dynamics of cellular metabolism in vivo. *Nat*. Biomed. Eng. 4, 335–342 (2020).

32. Uthayakumar, B. et al. Task activation results in regional^13^ C-lactate signal increase in the human brain. J. Cereb. Blood Flow Metab. 45, 1223–1231 (2025).

33. Liu, Y. et al. Interleaved fluid-attenuated inversion recovery (FLAIR) MRI and deuterium metabolic imaging (DMI) on human brain in vivo. Magn. Reson. Med. 88, 28–37 (2022).

34. Ahmadian, N. et al. Reproducibility of the Determination of^13^ C-Labeling of Glutamate and Glutamine in the Human Brain Using selPOCE-MRS at 7 T Upon [U-^13^ C]–Labeled Glucose Infusion. NMR Biomed. 38, e70026 (2025).

35. Schür, R. R. et al. Brain GABA levels across psychiatric disorders: A systematic literature review and meta-analysis of^1^ H-MRS studies. Hum. Brain Mapp. 37, 3337–3352 (2016).

36. Levy, L. M. & Degnan, A. J. GABA-based evaluation of neurologic conditions: MR spectroscopy. AJNR Am. J. Neuroradiol. 34, 259–265 (2013).

37. Cherix, A. et al. Metabolic signature in nucleus accumbens for anti-depressant-like effects of acetyl-L-carnitine. eLife 9, 1–19 (2020).

38. Hermans, L. et al. Brain GABA Levels Are Associated with Inhibitory Control Deficits in Older Adults. J. Neurosci. 38, 7844–7851 (2018).

39. Bell, T., Stokoe, M. & Harris, A. D. Macromolecule suppressed GABA levels show no relationship with age in a pediatric sample. Sci. Rep. 11, 722 (2021).

40. Hyder, F. Commentary to “Task activation results in regional^13^ C-lactate signal increase in the human brain”. J. Cereb. Blood Flow Metab. 45, 1413–1416 (2025).

41. Niess, F. et al. Noninvasive 3-Dimensional 1H-Magnetic Resonance Spectroscopic Imaging of Human Brain Glucose and Neurotransmitter Metabolism Using Deuterium Labeling at 3T: Feasibility and Interscanner Reproducibility. Invest. Radiol. 58, 431–437 (2023).

42. Otsuki, T. et al. Carbon 13–Labeled Magnetic Resonance Spectroscopy Observation of Cerebral Glucose Metabolism: Metabolism in MELAS: Case Report. Arch. Neurol. 62, 485 (2005).

43. Wijnen, J. P. et al. In vivo 13C magnetic resonance spectroscopy of a human brain tumor after application of 13C-1-enriched glucose. Magn. Reson. Imaging 28, 690–697 (2010).

44. Rothman, D. L. et al. Localized proton NMR observation of [3-^13^ C] lactate in stroke after [1-^13^ C] glucose infusion. Magn. Reson. Med. 21, 302–307 (1991).

45. Shulman, R. G., Hyder, F. & Rothman, D. L. Cerebral energetics and the glycogen shunt: Neurochemical basis of functional imaging. Proc. Natl. Acad. Sci. 98, 6417–6422 (2001).

46. Lizarbe, B. et al. Feasibility of in vivo measurement of glucose metabolism in the mouse hypothalamus by 1 H-[ 13 C] MRS at 14.1T. Magn. Reson. Med. 80, 874–884 (2018).

47. Xin, L., Lanz, B., Lei, H. & Gruetter, R. Assessment of metabolic fluxes in the mouse brain in vivo using 1 H-[13 C] NMR spectroscopy at 14.1 Tesla. J. Cereb. Blood Flow Metab. 35, 759–765 (2015).

48. Harris, A. D., Puts, N. A. J., Barker, P. B. & Edden, R. A. E. Spectral-editing measurements of GABA in the human brain with and without macromolecule suppression: Relationship of GABA+ and MM-suppressed GABA. Magn. Reson. Med. 74, 1523–1529 (2015).

49. Öz, G. et al. Advanced single voxel 1 H magnetic resonance spectroscopy techniques in humans: Experts’ consensus recommendations. NMR Biomed. 34, 1–18 (2021).

50. Andreychenko, A., Boer, V. O., Arteaga De Castro, C. S., Luijten, P. R. & Klomp, D. W. J. Efficient spectral editing at 7 T: GABA detection with MEGA-sLASER. Magn. Reson. Med. 68, 1018–1025 (2012).

51. Stefan, D. et al. Quantitation of magnetic resonance spectroscopy signals: the jMRUI software package. Meas. Sci. Technol. 20, 104035 (2009).

52. Cabanes, E., Confort-Gouny, S., Le Fur, Y., Simond, G. & Cozzone, P. J. Optimization of Residual Water Signal Removal by HLSVD on Simulated Short Echo Time Proton MR Spectra of the Human Brain. J. Magn. Reson. 150, 116–125 (2001).

53. Schwab, M., Bauer, R. & Zwiener, U. The distribution of normal brain water content in Wistar rats and its increase due to ischemia. Brain Res. 749, 82–87 (1997).

54. Xin, L., Lanz, B., Frenkel, H. & Gruetter, R. BISEP-based, Ultra-short TE 1 H–[ 13 C] NMR Spectroscopy of the Rat Brain at 14.1 T. www.jstage.jst.go.jp/browse/islsm (2009).

55. Cherix, A., Sonti, R., Lanz, B. & Lei, H. In Vivo Metabolism of [1,6-13C2]Glucose Reveals Distinct Neuroenergetic Functionality between Mouse Hippocampus and Hypothalamus. Metabolites 11, 50 (2021).

56. Shah, S., et al. Rapid fieldmap estimation for cardiac shimming. in Proc Intl Soc Mag Reson Med vol. 17 566 (2009).

57. Gruetter, R. & Tkac, I. Field mapping without reference scan using asymmetric echo-planar techniques. Magn. Reson. Med. 43, 319–323 (2000).

58. Klomp, D. W. J., Bitz, A. K., Heerschap, A. & Scheenen, T. W. J. Proton spectroscopic imaging of the human prostate at 7 T. NMR Biomed. 22, 495–501 (2009).

59. Tkac, I., Starcuk, Z., Choi, I.-Y. & Gruetter, R. In vivo1H NMR spectroscopy of rat brain at 1 ms echo time. Magn. Reson. Med. 41, 649–656 (1999).

60. Ratiney, H. et al. Time-domain semi-parametric estimation based on a metabolite basis set. NMR Biomed. 18, 1–13 (2005).

61. Ferrannini, E. et al. Effect of insulin on the distribution and disposition of glucose in man. J. Clin. Invest. 76, 357–364 (1985).

62. Krishnan, R. K., Evans, W. J. & Kirwan, J. P. Glucose Clearance Is Delayed after Hyperglycemia in Healthy Elderly Men. J. Nutr. 133, 2363–2366 (2003).

63. Gruetter, R., Seaquist, E. R. & Ugurbil, K. A mathematical model of compartmentalized neurotransmitter metabolism in the human brain. Am. J. Physiol.-Endocrinol. Metab. 281, E100–E112 (2001).

64. Shen, J. et al. Determination of the rate of the glutamate/glutamine cycle in the human brain by in vivo 13C NMR. Proc. Natl. Acad. Sci. U. S. A. 96, 8235–8240 (1999).

65. Lanz, B., Gruetter, R. & Duarte, J. M. N. Metabolic Flux and Compartmentation Analysis in the Brain In vivo. Front. Endocrinol. 4, 1–18 (2013).

66. Henry, P. G. et al. In vivo 13C NMR spectroscopy and metabolic modeling in the brain: a practical perspective. Magn. Reson. Imaging 24, 527–539 (2006).

