## Supplementary Figure legends and Supplementary material for "Non-invasive measurement of neurotransmitter-specific glucose metabolism in the human brain using proton-observed proton-edited ^13^C-MRS (POPE^13^C-MRS)"

Supplementary Methods

**POPE^13^C-MRS (Mouse)**

MRI/MRS acquisition: Offsets of the editing pulses were set to 1.9 ppm (editing ON) and 7.5 ppm (editing OFF) for GABA MEGA-sLASER and 4.1 ppm (editing ON) and 5.3 ppm (editing OFF) for lactate MEGA-sLASER with an editing pulse bandwidth of 130 Hz. Spectra were acquired with N=2048 points and a spectral width of 9.82 ppm. The repetition time (TR) was optimized for the GABA resonance and set to 3 s. In Study 1, the echo times (TE) were optimized for detection of ^13^C-satellites on 2 animals following a continuous 2h30min s.c. administration of [U-^13^C_6_]-Glc and were subsequently set to TE_GABA_=69.5ms and TE_LAC_=106ms for study 2, which included 5 male mice. Water suppression was performed using VAPOR^26^ at 4.7 ppm. Outer Volume Suppression (OVS) modules using hyperbolic secant pulses (4 mm thick slices and 0.5 mm gap) were interleaved with water suppression RF pulses. A baseline spectrum for GABA and lactate POPE^13^C-MRS was acquired prior to the ^13^C-glucose infusion and included 5 averages of 64 transients (32 x 2 editing condition) for each module leading to a 16min acquisition time. Thereafter, GABA and lactate POPE^13^C-MRS scans were acquired over 60 min in interleaved order with 64 transients (32 x 2 editing conditions; 3 min 12 s) repeated for each module. For quantification, a reference spectrum was acquired without water suppression after the baseline scan.

MRS data processing:

MRS spectral fitting was done using a homemade basis set used as prior knowledge for peak quantification in the time domain with QUEST, using a frequency range ($\pm$10Hz), damping range (10-20Hz) and phase range ($\pm$10deg) as fitting constraints. The basis set (Supplementary Fig.1) was obtained from phantom solutions of metabolites and included the following set of resonances: GABA^12^C-H3; GABA^12^C-H4; Gln^12^C-H2; Gln^13^C-H2; Glu^12^C-H2; Glu^13^C-H2; Gln^12^C-H4; Gln^13^C-H4; Glu^12^C-H4; Glu^13^C-H4; Gln^12^C-H3; Glu^12^C-H3 (the two satellite components of Gln^13^C-H4 and Glu^13^C-H4 were fitted as independent resonances, as their exact phases and resonance were not known *in vivo*). The satellite peaks were modelled using their heteronuclear J-coupling constant (^1^J_CH_) either determined experimentally in vivo (Glx(^13^C)H2 and Lac(^13^C)H3) or estimated from the literature. Cramer-Rao Lower Bounds (CRLB) were calculated by dividing the fit error (SD) obtained from QUEST quantification over the signal amplitude and multiplied by 100^29^.

Individual resonances were not corrected for differences in T2 decay or differences in editing efficiency between metabolites. The quantification values from the baseline spectrum (t_0_) and post-^13^C-glucose spectrum (t_f_) were corrected for potential motion or frequency drift artefacts using the tCr peak, assuming tCr signal is constant between t_0_ and t_f_. Concentration of ^13^C-labelled Glu was measured using the Glu(H4)_t0_-Glu(H4)_tf_ difference, which was then referenced to the water signal to get the labelled metabolite concentration. Similarly, the concentration of labelled GABA was measured using the GABA(H4)_t0_-GABA(H4)_tf_ difference and lactate using the Lac(H3)_t0_-Lac(H3)_tf_ difference normalized to the water reference. The Glx(^12^C)H2 and Glx(^13^C)H2 satellites resonances were used to estimate the isotopic fractional enrichment (FE) of GlxH2, reflecting overall labelling efficiency. FE was calculated as the ratio of the ^13^C-satellite signal to the total of GlxH2 signal, i.e. FE = Glx(^13^C)H2 / Glx(^12^C)H2 + Glx(^13^C)H2, under the assumption that the decrease in Glx(^12^C)H2 matches the increase in Glx(^13^C)H2. Metabolite labelling concentrations were also reported as a ratio to GlxH2_FE_ to correct the variation in ^13^C-labelling between subjects that may arise from differences in glucose infusion time or efficiency.

Supplementary Figures and Legends

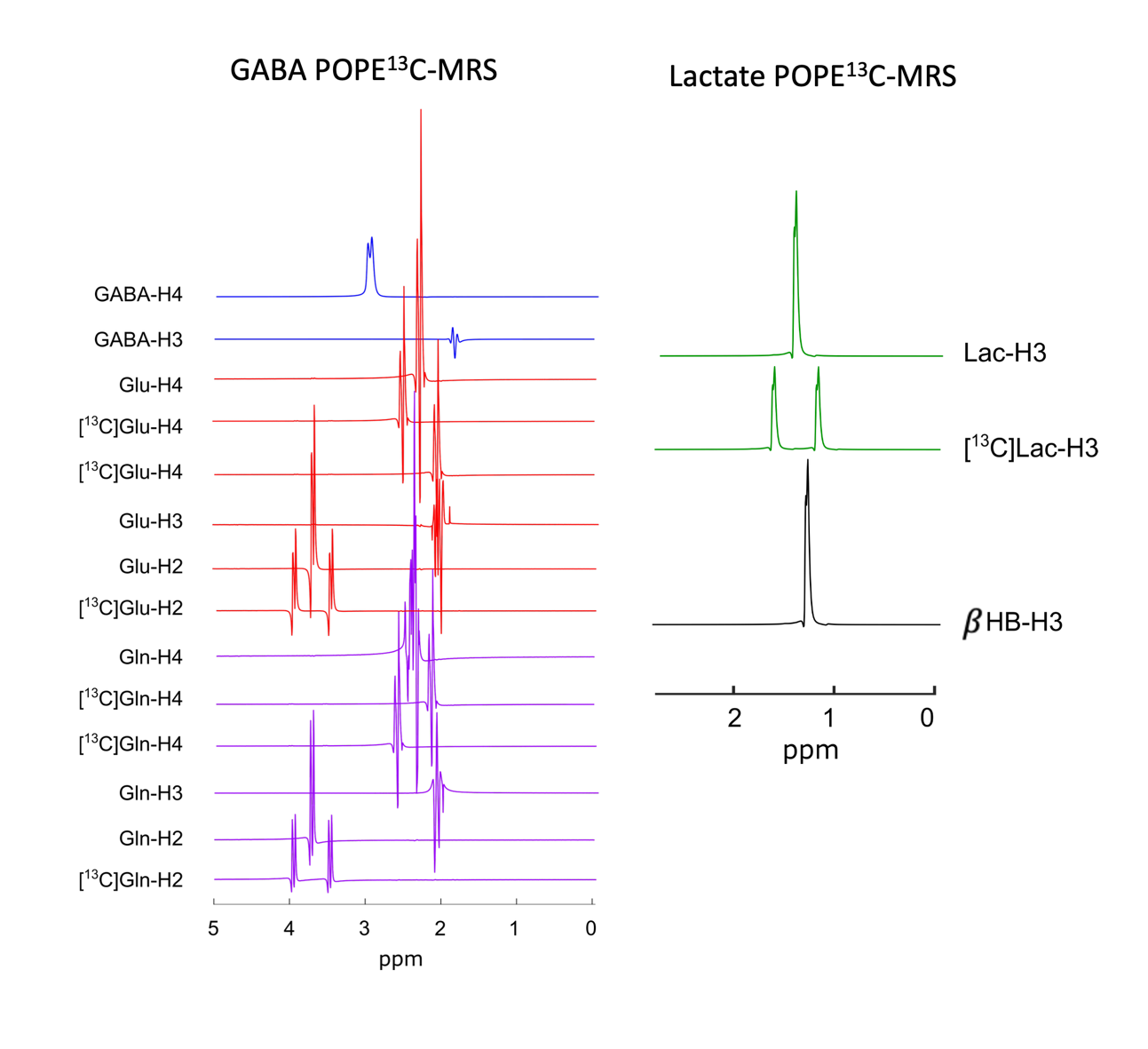

***Supplementary Figure 1: Basis set for spectral fitting in POPE^13^C-MRS***

*Individual fitting components of GABA POPE^13^C-MRS include (blue) GABA H3 and H4, (red) glutamate H2, H3, H4 with ^13^C-satellites for H2 and H4, (purple) glutamine H2, H3 and H4, with ^13^C-satellites for H2 and H4. Components of lactate POPE^13^C-MRS include (green) lactate H3 and H3 ^13^C-satellites and (black) beta-hydroxybutyrate (*$\beta$*HB) H3.*

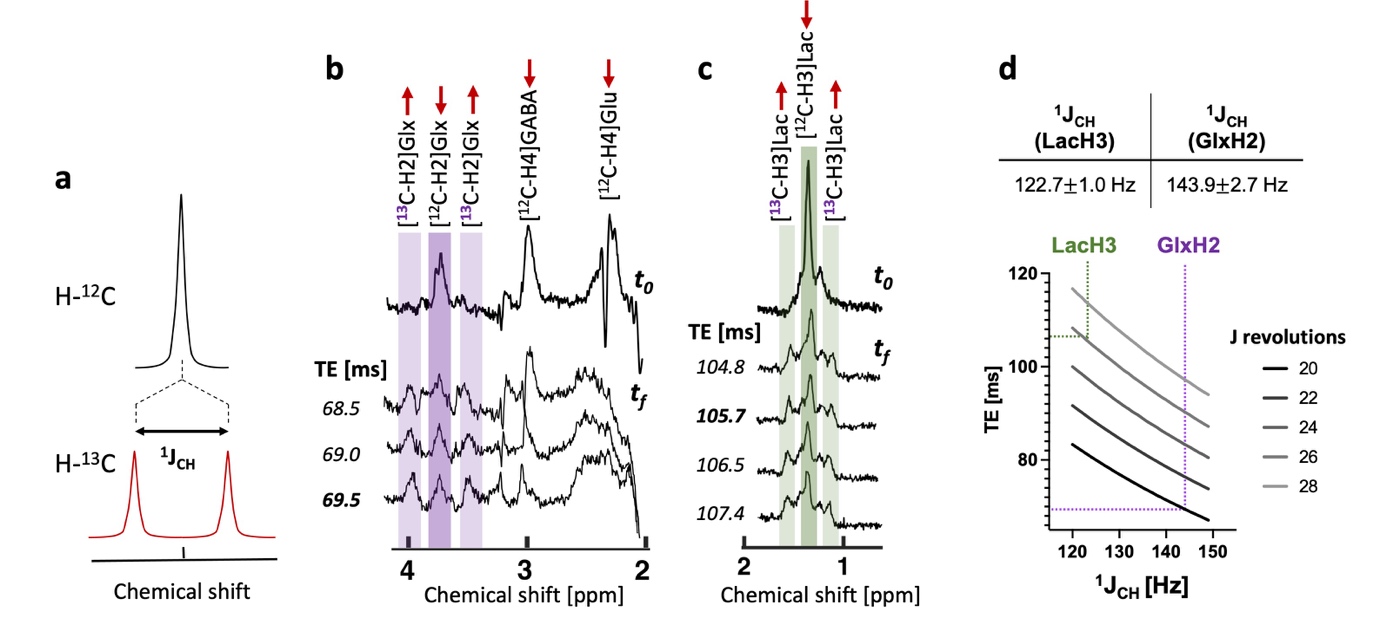

***Supplementary Figure 2: Heteronuclear J_CH_-coupling constants of Glx and lactate signals***

*(a) Schematic view of the heteronuclear J-coupling effects following ^13^C-labelling in POPE^13^C-MRS. Following the replacement of a ^12^C atom with a ^13^C atom on an adjacent (^1^J) proton, the signal splits in two equivalent signals of equal amplitude and separated by a frequency offset equal to the heteronuclear J-coupling constant (J_CH_). Satellite peaks identified (b) for GlxH2 in GABA-POPE^13^C-MRS and (c) for LacH3 in lactate-POPE^13^C-MRS after 120 min infusion of [U-^13^C_6_]-glucose. (d) Estimation of the echo time (TE) for optimal detection of GlxH2 and LacH3 satellite peaks at various number of J revolutions. Data shown as mean* $\pm$ *s.d., with s.d. being measured over all the different TEs. These experiments were performed in mouse brain under isoflurane anaesthesia.*

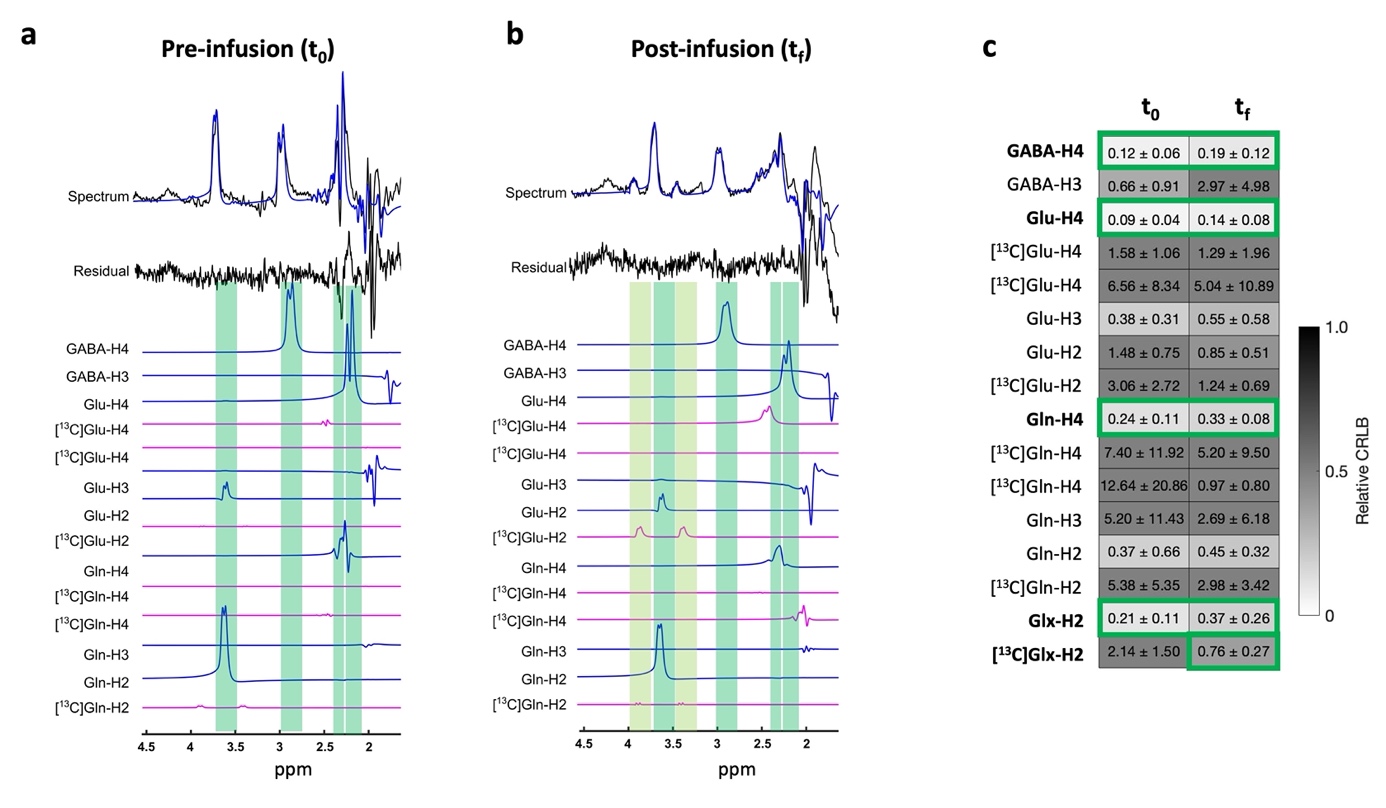

***Supplementary Figure 3: Metabolite quantification in POPE^13^C-MRS***

***a,b*** *Typical fitting result reproduced from QUEST analysis for (a) baseline spectrum and (b) spectrum of GABA POPE^13^C-MRS acquired after one hour of ^13^C-glucose infusion in the mouse brain. A 2 Hz Lorentzian apodization was applied for representation of the spectra, but not to the noise residual.* ***c*** *Relative quantification error (relative CRLB) for each component of the basis set before (t_0_) and after (t_f_) the infusion (n=5, mean*$\pm$*SD).*

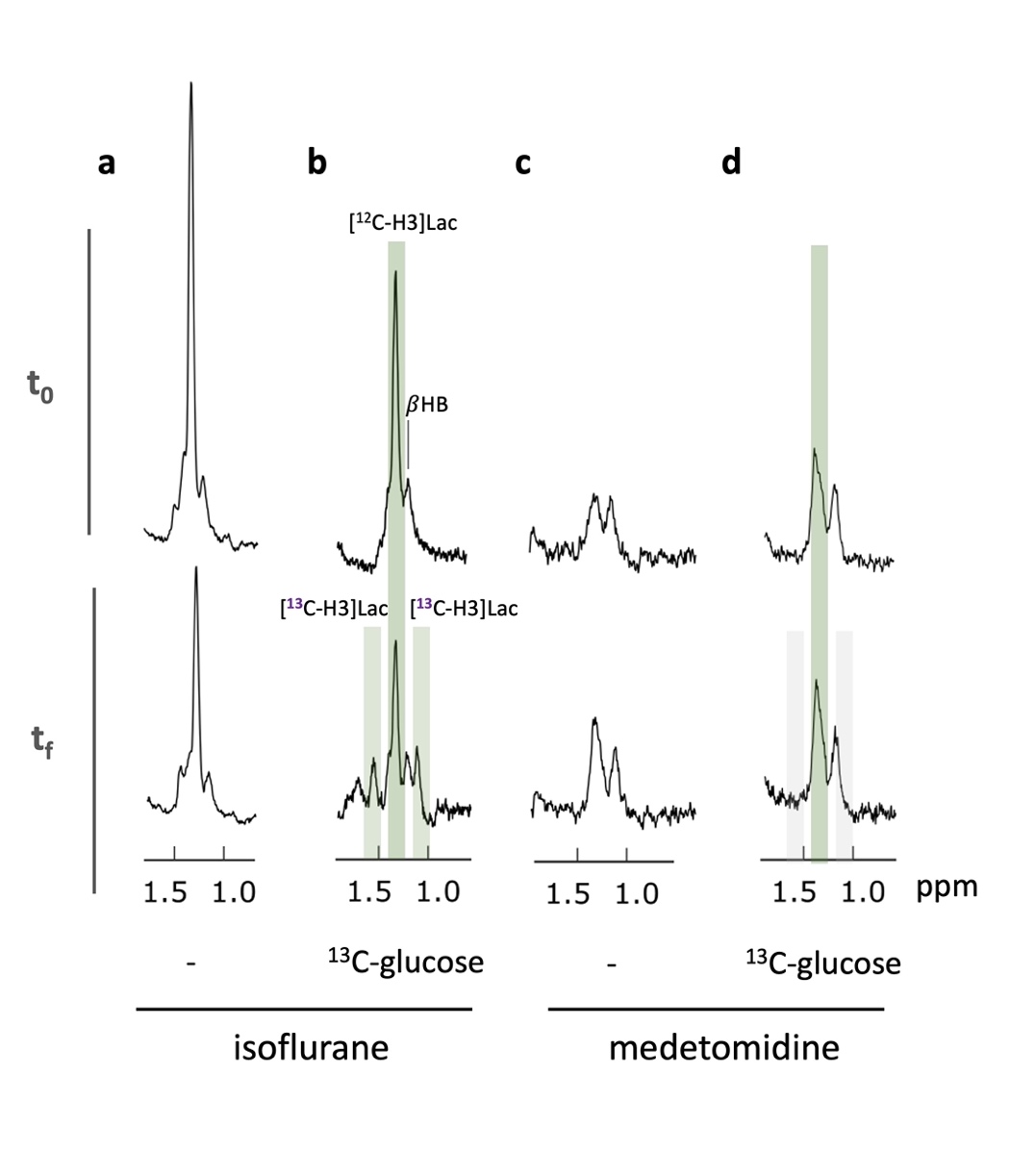

***Supplementary Figure 4: Detection of lactate ^13^C-satellite peaks under different anaesthetic regimen***

***a-d*** *Lactate signal acquired with lactate POPE^13^C-MRS in mouse brain before (t_0_) and after (t_f_) a one-hour infusion of [U-^13^C_6_]-glucose. Experiments were done under isoflurane with (b) and without (a) ^13^C-glucose infusion and under medetomidine with (d) and without (c) ^13^C-glucose infusion. Spectra are shown with 2Hz Lorentzian apodization.*

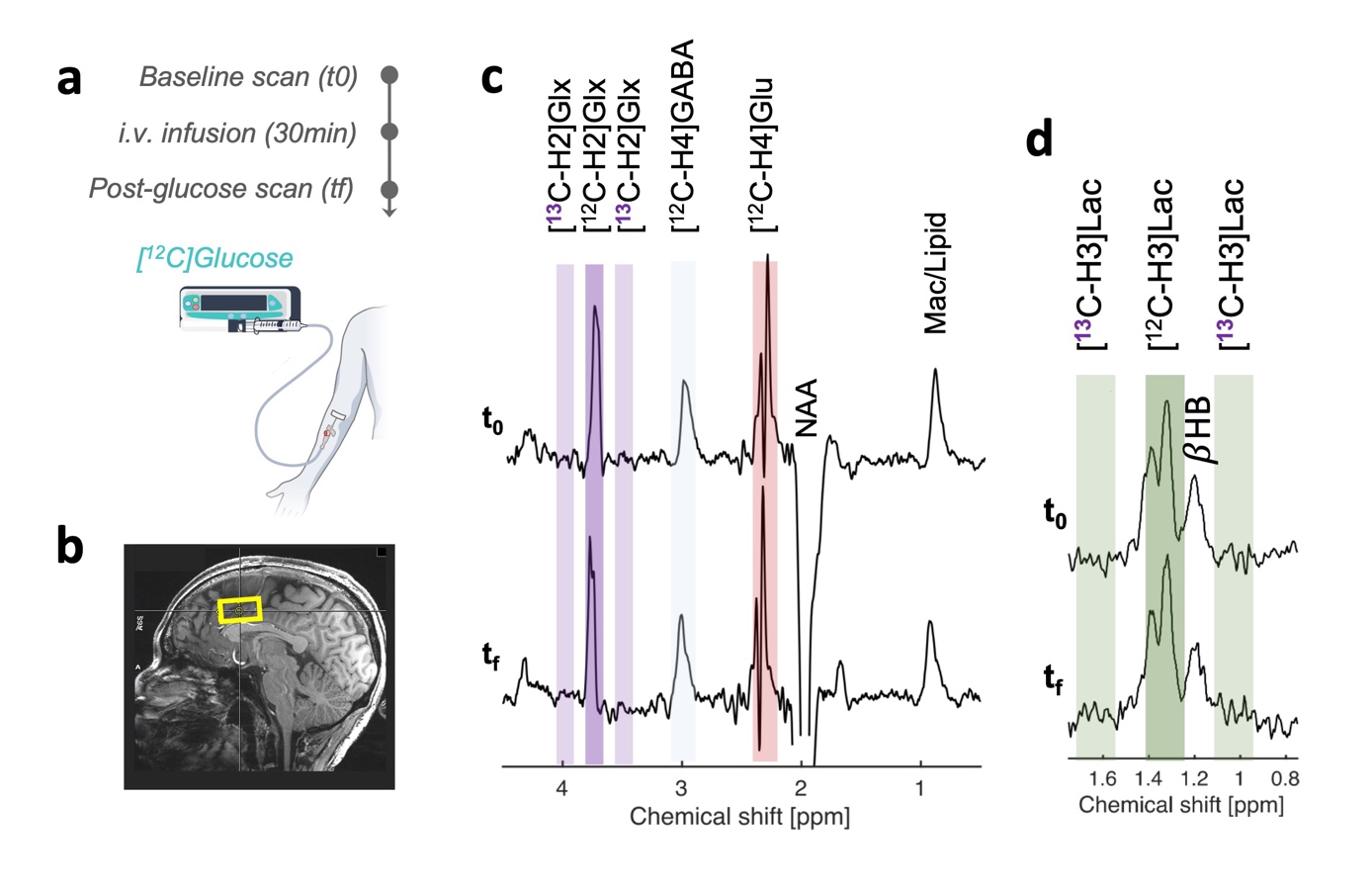

***Supplementary Figure 5: Stability of POPE^13^C-MRS upon glucose infusion***

***a*** *Timeline of the intravenous [^12^C]-glucose infusion and POPE^13^C-MRS baseline (t0) and post-infusion (tf) acquisitions.* ***b*** *Voxel positioning in dorsal anterior cingulate cortex (dACC) for POPE^13^C-MRS acquisition.* ***c-d*** *Average GABA-POPE^13^C-MRS (c) and lactate-POPE^13^C-MRS (d) spectra before (t0) and after (tf) the [^12^C]-glucose infusion. NAA, N-acetyl-aspartate;* $\beta$*HB, hydroxybutyrate; Mac/Lipid, macromolecules or lipids; Glx = glutamate + glutamine. Spectra are shown with 2Hz Lorentzian apodization.*

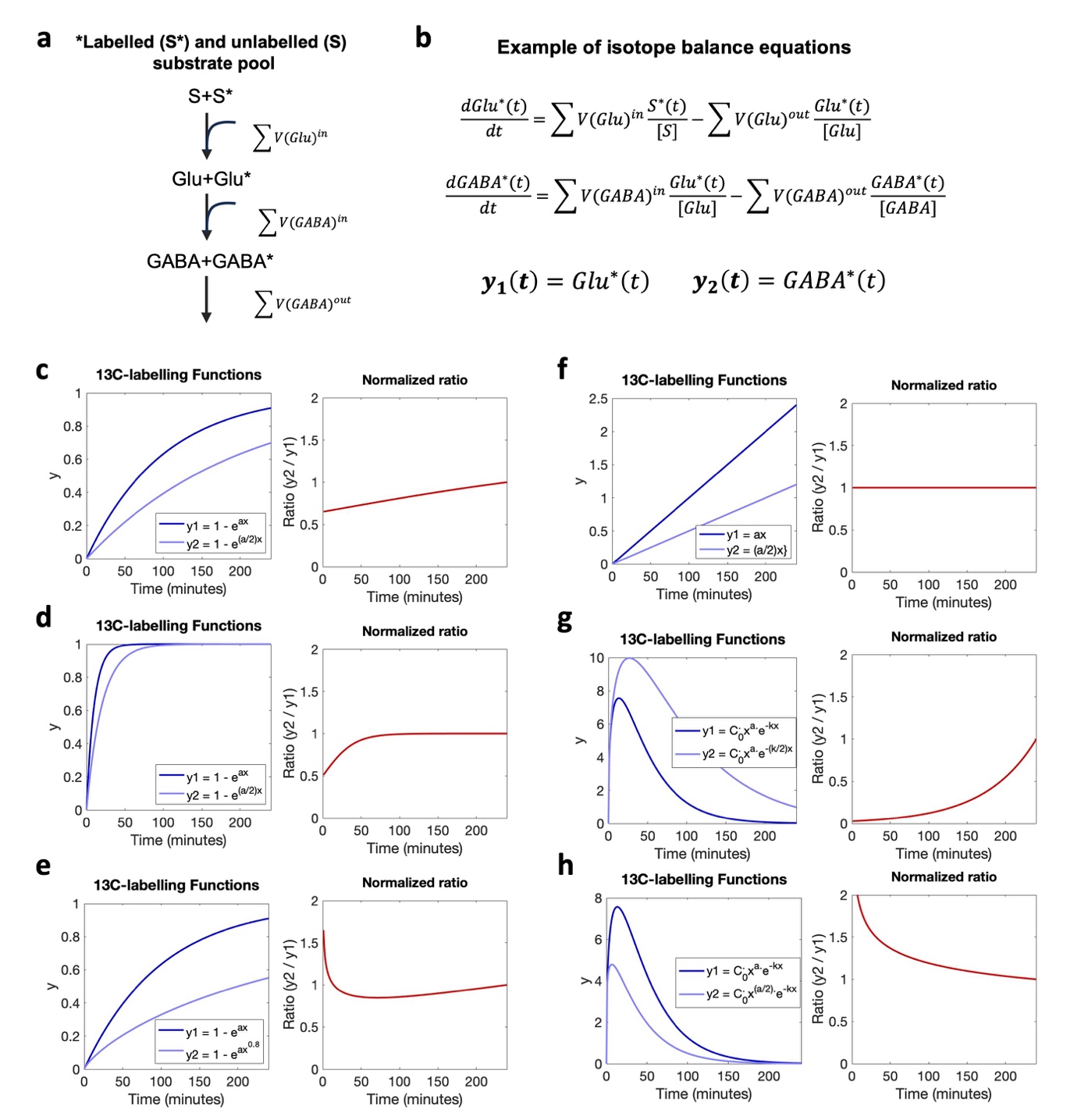

***Supplementary Figure 6: Example of labelling functions and associated ^13^C-labelling metabolite ratio stability with time***

***a,b*** *Example of kinetic modeling for two metabolite pools (e.g. GABA and Glutamate) labelled from a labelled substrate (S*). Labelling of each pool can be described with isotope balance equations and can lead to a variety of labelling functions y(t). For more details see*^65,66^*.*

***c-h*** *Example ^13^C-labelling functions that can be observed with various infusion protocols, such as continuous (c,d,e) or bolus (g,h) administration. Functions here were described as exponential saturation (y = 1-exp(a^.^x)) for high (c) or low a (d) values, (e) stretched exponential saturation (y = 1-exp(a^.^x^b^)), where b = 0.8, (f) linear (y = a^.^x) or (g,h) gamma-variate function (y = C_0_^.^x^a.^exp(-kx)). Note that all labelling functions here aren’t solutions of the isotope balance equation presented in b (e.g. linear function) but represent potential labelling profiles shown as examples.*

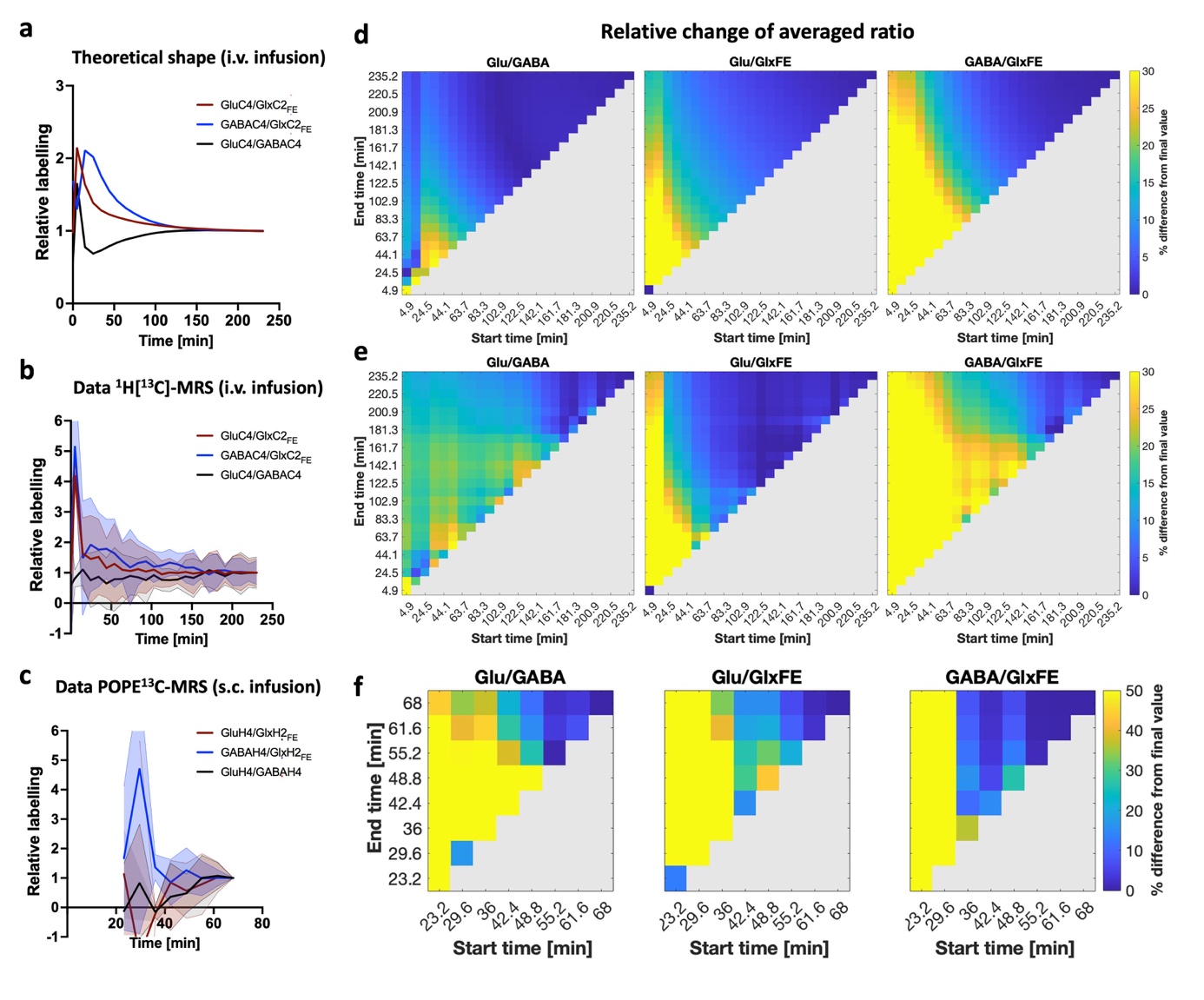

***Supplementary Figure 7: Time and acquisition window dependencies of metabolite labelling ratios in mice***

***a,b,c*** *Time effect and stability of ^13^C-labelled metabolite ratios established from a previous study^30^ (a,b) or in the current study with POPE^13^C-MRS (c) in mice. (a) the theoretical labelling values inferred from a pseudo-3 compartment model or (b) the associated raw data. The data was acquired in mouse brain following intravenous (a,b) or subcutaneous (c) infusion of [U-^13^C_6_]-glucose. The shaded areas (b,c) represent the s.d.* ***d,e,f*** *Stability of ^13^C-labelled metabolite ratios for various data acquisition windows relative to the start and end of the ^13^C-glucose infusion, estimated from (d) the theoretical pseudo-3 compartment model values, (e) its associated raw data or from POPE^13^C-MRS (f) in mice.*
